# Microglia control glutamatergic synapses in the adult mouse hippocampus

**DOI:** 10.1101/2021.02.01.429096

**Authors:** Bernadette Basilico, Laura Ferrucci, Patrizia Ratano, Maria T. Golia, Alfonso Grimaldi, Maria Rosito, Valentina Ferretti, Ingrid Reverte, Maria C. Marrone, Maria Giubettini, Valeria De Turris, Debora Salerno, Stefano Garofalo, Marie-Kim St-Pierre, Micael Carrier, Massimiliano Renzi, Francesca Pagani, Marcello Raspa, Ferdinando Scavizzi, Cornelius T. Gross, Silvia Marinelli, Marie E. Tremblay, Daniele Caprioli, Laura Maggi, Cristina Limatola, Silvia Di Angelantonio, Davide Ragozzino

**Affiliations:** Department of Physiology and Pharmacology, Sapienza University of Rome, Rome, Italy; IRCCS Neuromed, Via Atinese 18, 86077, Pozzilli, IS, Italy; Center for Life Nanoscience, Istituto Italiano di Tecnologia, Rome, Italy; Dipartimento di Biologia e Biotecnologie “Charles Darwin”, Sapienza University of Rome, Rome, Italy; Santa Lucia Foundation (IRCCS Fondazione Santa Lucia), Rome, Italy; European Brain Research Institute-Rita Levi Montalcini, Rome, Italy; CrestOptics S.p.A., Via di Torre Rossa 66, 00165 Rome, Italy; Centre de Recherche du CHU de Québec, Axe Neurosciences Québec, QC, Canada; Département de médecine moléculaire, Université Laval Québec, QC, Canada; National Research Council, Institute of Biochemistry and Cell Biology (CNR-IBBC/EMMA/Infrafrontier/IMPC), International Campus “A. Buzzati-Traverso”, Monterotondo (Rome) Italy; Epigenetics and Neurobiology Unit, European Molecular Biology Laboratory (EMBL), Monterotondo, Italy; Division of Medical Sciences, University of Victoria, Victoria, Canada

**Keywords:** glutamatergic transmission, hippocampus, learning, microglia, neuron-microglia interaction

## Abstract

Microglial cells are active players in regulating synaptic development and plasticity in the brain. However, how these cells influence the normal functioning of synapses is largely unknown. In this study, we characterized the effects of pharmacological depletion of microglia, achieved by administration of PLX5622, on hippocampal CA3-CA1 synapses of adult wild type mice. Following microglial depletion, we observed a reduction of spontaneous and evoked glutamatergic activity associated with a decrease of dendritic spine density. We also observed the appearance of immature synaptic features accompanied by higher levels of plasticity. In addition, microglia depleted mice showed a deficit in the acquisition of the Novel Object Recognition task. Remarkably, microglial repopulation after PLX5622 withdrawal was associated with the recovery of hippocampal synapses and learning functions. Altogether, these data demonstrate that microglia contribute to normal synaptic functioning in the adult brain and that their removal induces reversible changes in synaptic organization and activity of glutamatergic synapses.

## INTRODUCTION

Microglia are the resident macrophage cells of the central nervous system (CNS) and their inflammatory role is well characterized (Rock *et al*, 2004). Beyond this, microglia perform a variety of other homeostatic functions and in particular support the development and maturation of neural circuits (Tremblay *et al*, 2010; Paolicelli *et al*, 2011; Schafer *et al*, 2012).

Studies in defective-microglia models, such as the *Cx3cr1* knockout (KO) mice (Paolicelli *et al*, 2014) have shown that the disruption of neuron-microglia signaling causes alterations in brain circuit development, resulting in reduced connectivity between brain areas (Zhan *et al*, 2014) and consequent impairment of cognitive functions (Rogers *et al*, 2011).

These alterations associated with a delay in synaptic maturation (Paolicelli *et al*, 2011; Hoshiko *et al*, 2012) have permanent consequences on synaptic properties (Basilico *et al*, 2019; Zhan *et al*, 2014) and plasticity (Maggi *et al*, 2011; Rogers *et al*, 2011). Notably, defective neuron-microglia signaling in the hippocampus profoundly affects the strength and reliability of glutamatergic synapses (Basilico *et al*, 2019). All these effects are ascribed to the ability of microglia during development to foster synaptic pruning (Paolicelli *et al*, 2011), likely by contacting and phagocyting synaptic elements (Tremblay *et al*, 2010; Schafer *et al*, 2012; Weinhard *et al*, 2018).

In the last decade, several studies started to unravel the constitutive role of microglia in regulating synaptic plasticity and remodeling of neuronal circuits during adulthood. Loss-of-function mutation or deletion studies of microglial DAP12, CD200R or CX3CR1 resulted in reduced or enhanced hippocampal long-term potentiation (LTP) (Roumier *et al*, 2004, 2008; Maggi *et al*, 2011; Rogers *et al*, 2011). These, along with other evidences (reviewed in (Paolicelli *et al*, 2014)), suggest that microglia might play a critical role in learning and memory processes during adulthood. Indeed, signaling deficiency in *Cx3cr1* knockout mice, microglial-BDNF deletion or microglial depletion resulted in impaired motor learning and deficits in fear conditioning, Morris water maze and Novel Object Recognition (NOR) tasks (Maggi *et al*, 2011; Rogers *et al*, 2011; Parkhurst *et al*, 2013).

In this context, and given the increasing evidence of microglia role in pathological processes (Marinelli *et al*, 2019), it is pivotal to understand how microglial activity regulates learning processes and synaptic functions in physiological conditions. To this aim, using chronic pharmacological inhibition of colony-stimulating factor 1 receptor (CSF1R) by systemic PLX5622 treatment (Dagher *et al*, 2015), we investigated the impact of microglial depletion and repopulation on structural and functional neural plasticity in the hippocampus, and its consequences on the acquisition of the NOR Task.

## RESULTS

### PLX5622 treatment causes microglial depletion in the hippocampus without triggering neuroinflammation

In order to decipher the role of microglia in adult hippocampal synaptic functioning, 5-weeks-old C57BL6/J mice were fed with a chow containing 1200 PPM of PLX5622 for 7 days (Fig. 1A).

**Figure 1.**
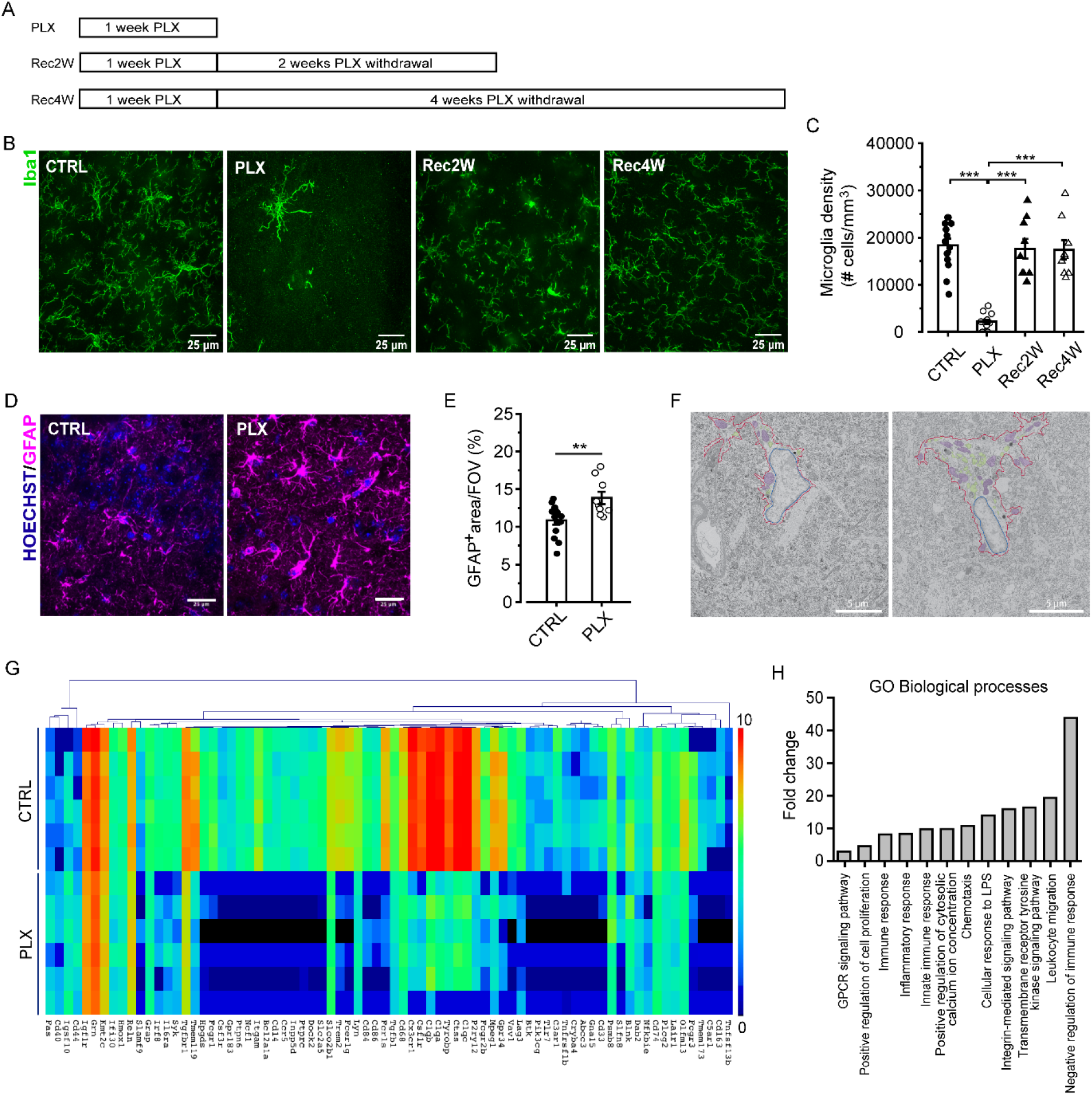
CSF1R inhibition leads to microglia depletion without causing neuroinflammation. A) Experimental design to deplete microglia. C57BL6/J mice are fed for 7 days with 1200 mg/kg PLX5622. The mice are then returned to standard chow and microglial density is assessed after 2 (Rec2W) and 4 weeks (Rec4W). B) Representative confocal images of Iba1+ microglial cells in the hippocampal CA1 *stratum radiatum*. C) Quantification of microglial density in the hippocampal CA1 *stratum radiatum* in control condition (n=13 fields/3 mice), after PLX5622 treatment (n=14 fields/4 mice), 2 weeks (n=9 fields/3 mice) and 4 weeks (n=9 fields/3 mice) after PLX5622 withdrawal. One-way ANOVA: F(3,44)=33.96, p<.001; Holm-Sidak post-hoc: ***p<.001. D) Representative images of astrocytes identified in hippocampal slices from control and PLX-treated mice immunolabeled with anti-GFAP antibody (magenta) and Hoechst for nuclei visualization (blue). 60x objective (scale bar = 25 µm) acquisition of the *stratum radiatum*. On the right, quantification of GFAP signal representing the area occupied by fluorescent GFAP+ cells *vs*. total field of view revealed increased astrogliosis in PLX-treated mice (n=34/9/2 fields/slices/mice) compared to control (n=54/14/3). t-test: t=-3.08, **p<.01. E) Electron micrograph of astrocytes in the *stratum radiatum* of the hippocampal CA1 in control versus PLX-treated groups using scanning electron microscopy. Red line = cytoplasm, Blue line = nucleus, purple = mitochondria, green = phagocytic elements. Scale bar = 5 µm. F) Heat map of unsupervised hierarchical clustering of the 76 differentially expressed genes in control and PLX hippocampus samples (n=6) analyzed by Nanostring (72 genes downregulated and 4 up-regulated in PLX samples). Colors in the heatmap indicate log2 counts normalized to housekeeping genes. G) Gene Ontology enrichment analysis in ‘Biological Process’ categories for 72 genes down-regulated in PLX samples and identified according to Database for Annotation, Visualization and Integrated Discovery (DAVID) functional annotation. Data presented as Mean ± SEM (C, E) or in Log2 scale (H).

As previously reported (Huang *et al*, 2018; Zhan *et al*, 2019), PLX-treated mice showed a 90% reduction of Iba1+ cells in the CA1 *stratum radiatum* of the hippocampus relative to control (Fig. 1B,C). After discontinuing the PLX5622 diet for two weeks, microglia repopulated the mouse brain (Fig. 1B,C).

Microglial depletion may cause profound changes in the brain parenchyma due to a potential alteration of tissue homeostasis or to the death of microglia. To determine whether the effects of PLX on synapses result from an alteration of the brain inflammatory state, we performed immunofluorescence staining of astrocytic and inflammatory markers. We observed an increase of GFAP+ signal density in CA1 *stratum radiatum*, indicative of reactive astrogliosis in microglia-depleted mice (Fig. 1D,E). In the ultrastructural analysis of astrocyte in electron micrograph of microglia-depleted mice, we observed a very active cytoplasm, characterized by a higher number of phagocytic vesicles in an overall larger size, typical of their important phagocytic activity (Fig. 1F). Besides the astrocyte’s active cytoplasm, we counted numerous mitochondria, indicating a high-energy need (Fig. 1F). Conversely, we did not observe clear ultrastructural signs of neuroinflammation in microglia-depleted mice, as revealed by the absence of organellar disruption throughout the neuropil in both PLX-treated and control mice (Fig. EV1). Qualitative analysis of the whole neuropil of the *stratum radiatum* did not reveal the presence of stressed cells in PLX-treated mice, as revealed by electron-dense cyto- and nucleoplasm. In addition, we did not observe the presence of dark microglia in the *stratum radiatum* of both PLX and control mice, supporting the absence of inflammatory processes. Moreover, we did not identify blatant ultrastructural signs of cell death in microglia-depleted mice using scanning electron microscopy.

To further confirm the absence of inflammation, we performed gene expression profiling of the hippocampus (Nanostring analysis of total hippocampal RNA extracts). We observed changes in gene expression for 76 over 757 genes within the Neuroinflammation mouse panel in microglial-depleted mice (Fig. 1G). Among these, 72 were downregulated in PLX-treated mice, while only 4 (FAS, CD44, CD40, IGSF10) were upregulated. The 72 downregulated genes belong to microglia-mediated inflammatory function as determined by Gene ontology and Kegg-pathway analysis, as for instance genes related to complement cascade (C1QA, C3AR1, C1QB, C1QC; p<0.001), microglial G-protein signaling pathway (P2RY12, CCR5, CX3CR1), and natural killer cell cytotoxicity (VAV1, PIK3CG, PTPN6, SYK; p<0.001) (Fig. 1H), corroborating the absence of an inflammatory state upon PLX treatment.

### Microglial depletion impairs glutamatergic synaptic activity in the hippocampus

To investigate whether microglial depletion could interfere with synaptic functional properties, we characterized hippocampal synapses in acute brain slices by patch clamp recordings in CA1 pyramidal neurons. The main finding was that PLX-treated mice showed a significant decrease in the amplitude of spontaneous excitatory postsynaptic currents (sEPSC) relative to control mice (Fig. 2A-D), without major effects on sEPSC frequency (Fig. 2E-F). Notably, this effect was specific for glutamatergic synapses, since we did not observe differences in amplitude (Fig. S1A) or frequency (Fig. S1B) of spontaneous GABAergic IPSCs.

**Figure 2.**
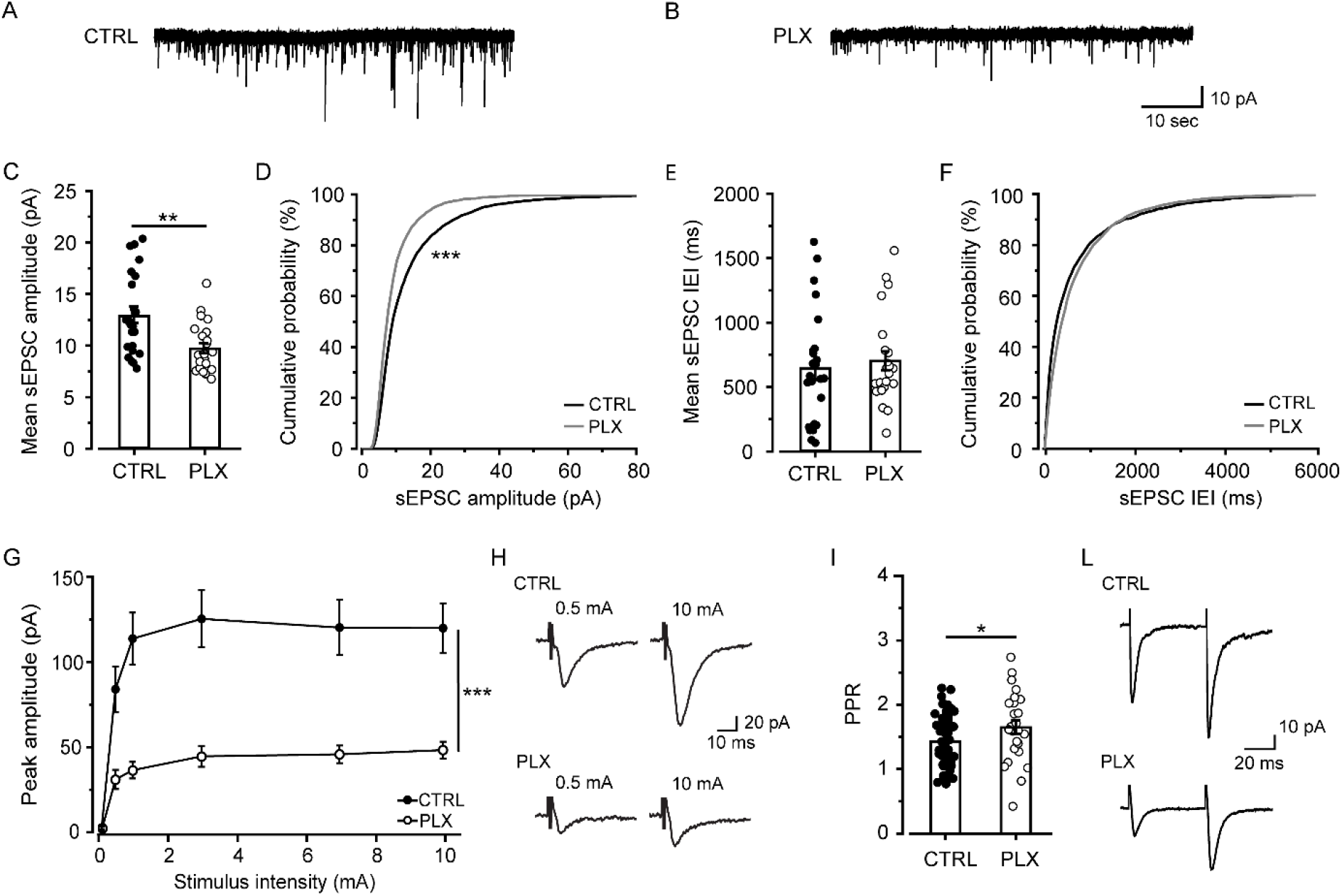
Microglia depletion impairs glutamatergic transmission in hippocampus. A-B) Representative traces of sEPSC recorded at -70 mV from CA1 pyramidal neurons in control (A) and PLX-treated mice (B). C) Bar graph of mean sEPSC amplitudes in control (n=25 cells/6 mice) and PLX neurons (n=23 cells/9 mice). t-test: t=3.22, **p<.01. D) Cumulative distributions of sEPSC amplitudes as in (C). K-S test: D=.15, ***p<.001. E) Bar graph of mean sEPSC interevent interval in control (n=25 cells/6 mice) and PLX neurons (n=23 cells/9 mice). F) Cumulative distributions of sEPSC interevent interval as in (E). G) Input-output curve of evoked EPSC peak amplitudes recorded at −70 mV from control (n=24 cells/8 mice) and PLX neurons (n=24 cells/7 mice). Note that in PLX-treated mice, neurons show significantly lower peak amplitudes compared to control. Two-way ANOVA: treatment F(1,275)=105.06 ***p<.001; stimulation F=(5,275)=20.46 p<.001; interaction F(5,275)=4.44 p<.001. H) Representative EPSCs recorded at −70 mV from control and PLX neurons, following Schaffer collateral stimulation at 0.5 mA and 10 mA, respectively. I) Bar graph showing the PPR, determined by dividing the second EPSC by the first, recorded in control (n=58 cells/20 mice) and PLX-treated mice (n=26 cells/10 mice). t-test: t=-2.16, *p<.05. L) Representative eEPSC induced by paired-pulse stimulation of Schaffer collateral. Data presented as Mean ± SEM.

To further investigate the impact of microglial depletion on hippocampal synaptic transmission, we analyzed evoked EPSC at CA3-CA1 synapses (Basilico *et al*, 2019). As revealed by the input/output curve, EPSCs recorded at hyperpolarized potential (-70 mV; AMPA component) displayed strongly reduced amplitudes in PLX-treated mice relative to controls (Fig. 2G-H), suggesting that microglial depletion profoundly affects CA3-CA1 synaptic functional connectivity. Notably, this effect was not observed when experiments were repeated in *Cx3cr1*^-/-^ mice (mice lacking the microglial fractalkine receptor), indicating that PLX-induced synaptic dysfunction is directly correlated to functional neuron-microglia crosstalk (Fig. EV2). The reduction in EPSC amplitudes was not restricted to the AMPA component. Indeed, when we analyzed the NMDA receptor (NMDAR) component (holding cells at +40 mV membrane potential in the presence of NBQX 10 μM), we observed a reduction of current amplitude in PLX-treated mice (Fig. S2).

To better comprehend the origin of the deficient glutamatergic synaptic transmission observed in PLX-treated mice, we analyzed functional and structural parameters underlying synaptic efficacy (refer to next section).

### Microglial depletion causes a regression in the properties of CA1 synapses and enhances long-term potentiation

Current clamp analysis of CA3 neurons (stemming excitatory inputs onto CA1 pyramidal cells) showed that both cell excitability (Fig. S3A-B) and passive properties (Fig. S3C-E) were unaltered in PLX-treated mice, suggesting that microglial elimination impaired the CA3-CA1 glutamatergic transmission at the synaptic level.

We thus examined more in detail the AMPAR-mediated component of EPSCs, evoked in CA1 pyramidal neurons by stimulation of Schaffer collaterals. We observed a higher paired pulse ratio of EPSCs in slices from PLX-treated mice relative to controls (Fig. 2I-L), suggesting a reduction in the probability of glutamate release (Dobrunz & Stevens, 1997). Consistently, we did not observe a reduction in mEPSC amplitude (CTRL: 7.7 ± 0.4 pA; PLX: 8.6 ± 0.6; t-test, p=.37) or frequency (CTRL: 1.4 ± 0.7 Hz; PLX: 1.8 ± 0.3 Hz; t-test, p=.61) in slices from PLX-treated mice (not shown), pointing to a presynaptic origin of the synaptic defect. Interestingly, when experiments were repeated in conditions of high release probability (Ca^2+^/Mg^2+^ ratio = 8) (Basilico *et al*, 2019), the EPSC amplitudes in the I/O curve were significantly lower in slices from PLX-treated animals compared to control ones (n= 9 CTRL vs n=7 PLX; Two-way ANOVA, p<.001; not shown).

Further, and remarkably, in patch clamp recordings from PLX-treated mice, we observed a reduction of synaptic multiplicity (indicated by the lack of difference between miniature and spontaneous EPSC amplitudes; Fig. 3A-D) (Hsia *et al*, 1998), as well as a reduction in the AMPA/NMDA ratio (Fig. 3E-F). Overall, these data suggest that microglia depletion is associated with the appearance of immature synaptic features (Hoshiko *et al*, 2012; Basilico *et al*, 2019).

**Figure 3.**
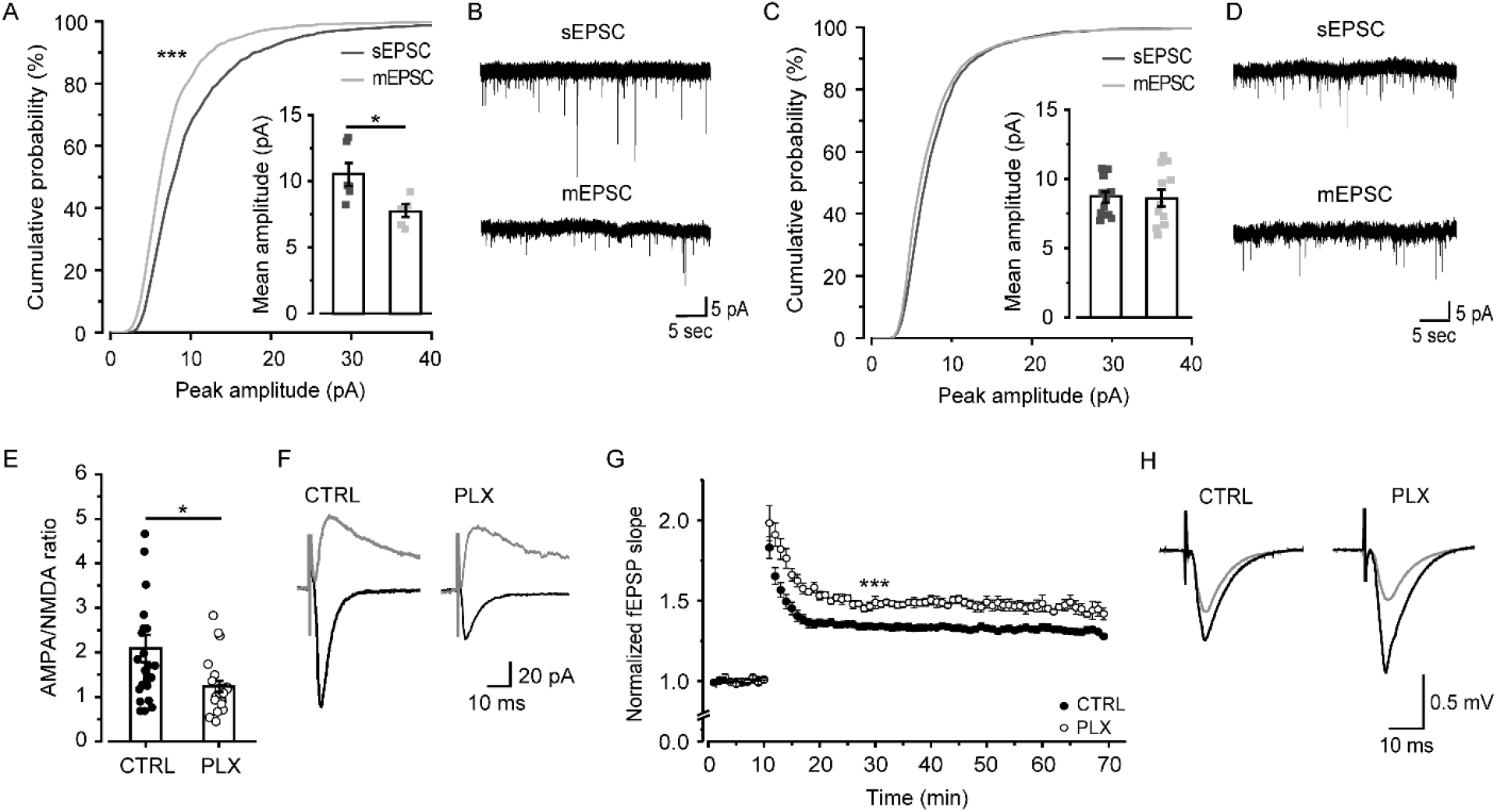
CA3-CA1 synapses show immature properties after microglia depletion. A) Cumulative distribution and scatter plot showing a significant difference between mEPSC and sEPSC amplitudes in control mice (n=6 cells/3 mice). paired t-test: t=3.47, *p<.05. K-S test: D=.22, ***p<.001. B) Representative sEPSC and mEPSC traces recorded at -70 mV from pyramidal neurons of control mice. C) In PLX-treated mice (n=12 cells/8 mice) the difference seen in CTRL mice for sEPSC and mEPSC amplitudes was lost (cumulative distribution and scatter plot), indicating a defect in synaptic multiplicity. D) Representative sEPSC and mEPSC traces recorded at -70 mV from pyramidal neurons of PLX mice. E) The AMPA/NMDA ratio is significantly lower in PLX (n=23 cells/10 mice) compared to control neurons (n=26 cells/10 mice) t-test: t=-2.47, *p<.05. F) Representative traces of AMPA-mediated currents, elicited at -70 mV, and NMDA-mediated currents (+40 mV), evoked by Schaffer collateral stimulation. G) Time course of fEPSP slope responses evoked at 0.05 Hz and normalized. Note that fEPSP amplitudes are higher in PLX-treated mice (CTRL, n=14 slices/6 mice; PLX, n= 8 slices/4 mice). t-test: t=-6.79, ***p<.001. H) Representative field potential waveforms before (gray) and after (black) 30 min of HFS induction, for each condition as indicated. Data presented as Mean ± SEM.

We then evaluated the effects of microglia depletion on synaptic plasticity in the hippocampal CA1 region applying a100 Hz stimulation of Schaffer collateral inputs. We observed a significant increase in LTP amplitude at CA1 synapses in PLX-treated mice (1.5 ± 0.03) relative to controls (1.3 ± 0.01) (Fig. 3G-H).

### Microglial depletion reduces spine density but did not alter spine morphology in the hippocampus

In order to investigate the structural determinant of the physiological changes described above, we analyzed the density, morphology and distribution of dendritic spines in the CA1 *stratum radiatum*. We observed that microglial depletion induces a significant reduction in spine density (Fig. 4A-B) without changes in spine morphology (Fig. 4C). Moreover, we observed a significant reduction of the postsynaptic marker drebrin in microglia-depleted mice (Fig. 4D-E), with no significant changes in synapsin I (presynaptic marker; Fig. 4F).

**Figure 4.**
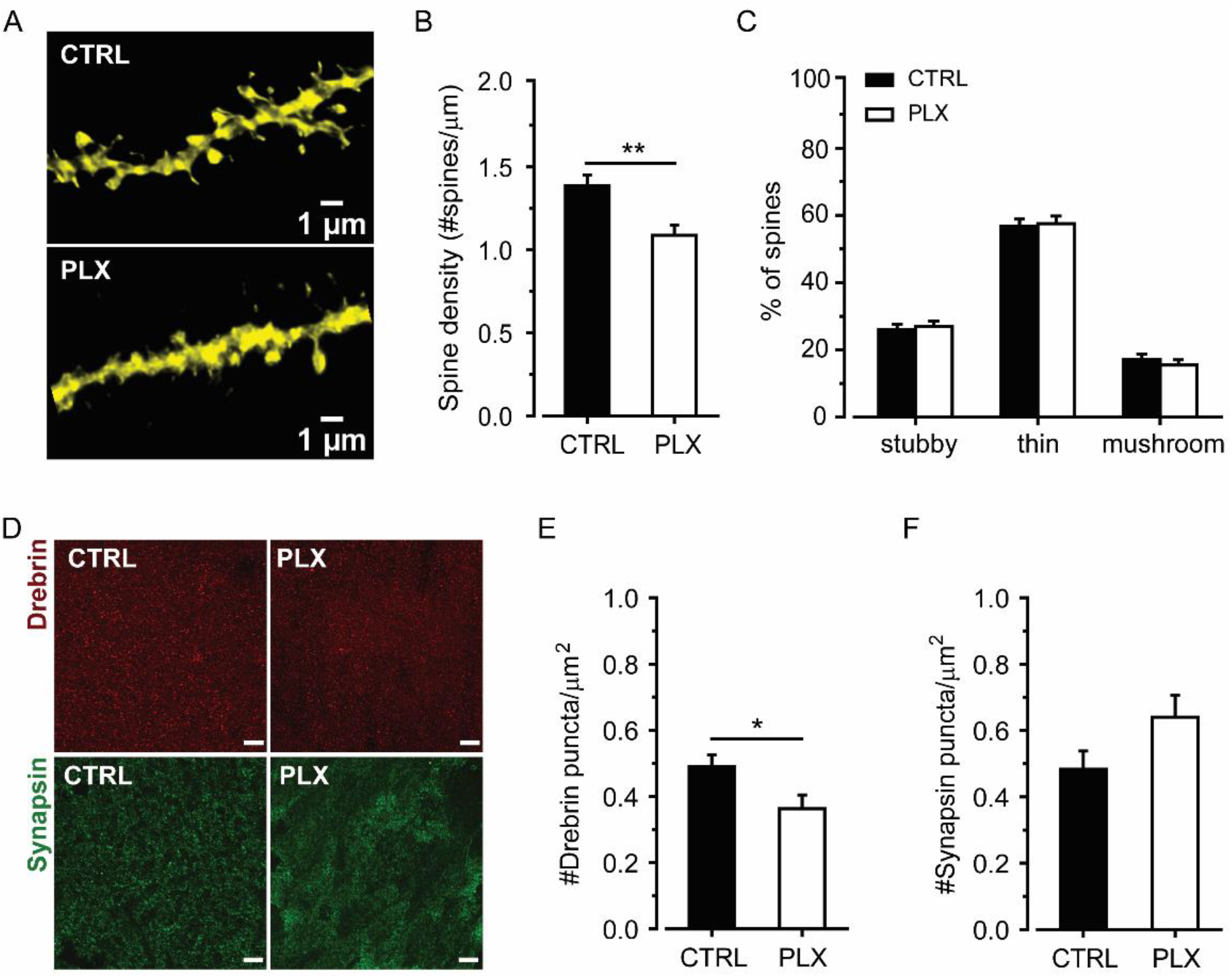
Microglia depletion impairs spine density in hippocampus. A) Representative confocal images of dendritic segments belonging to control and PLX pyramidal neurons from CA1 *stratum radiatum*. B) Bar graph showing the reduction of the overall spine density in PLX neurons (n=46 dendrites/5 mice; control n=46 dendrites/5 mice). t-test: t=3.28, **p<.01. C) Quantification of the percentage of stubby, thin and mushroom spines does not reveal any difference between the two experimental conditions. D) Super-resolution images of drebrin (red; left) and synapsin (green; right) synaptic signals in CA1 *stratum radiatum* of control and PLX-treated mice. E-F) Quantification of drebrin (E) (control n=18 fields/3 mice; PLX n=16 fields/3 mice) and synapsin (F) (control n=18 fields/3 mice; PLX n=17 fields/3 mice) signal density. t-test: *p<.05. Data presented as Mean ± SEM.

### Microglial repopulation is followed by recovery of synaptic function

It is well established that microglia replenish the brain once the PLX5622 diet is discontinued (Dagher *et al*, 2015; Zhan *et al*, 2019). However, these cells may present distinct morphological features compared to the normal microglia (Elmore *et al*, 2015). To address this question, we performed a qualitative ultrastructural analysis of microglia at 2 and 4 weeks after PLX5622 withdrawal. We observed that after 2 weeks of recovery from PLX diet (Rec2W) microglial cells display more mitochondria and phagocytic inclusions compared to untreated mice (Fig. 5A). However, after 4 weeks of PLX withdrawal (Rec4W) microglia display the typical cytoplasmic appearance (Fig. 5A). In line with these observations, our immunofluorescence analysis show that the repopulation process was associated with a progressive return to normal microglial morphology in the hippocampus (Fig. 5B-E). Indeed, after 2 weeks of PLX withdrawal (Rec2W) microglial density was fully restored (see Fig. 1B-C), but the cells showed a larger soma area and reduced ramification relative to untreated mice. After 4 weeks of withdrawal (Rec4W), microglial soma size returned to control levels but the number of branches was still low compared to untreated mice. Notably, this progressive recovery of microglia density and morphology was mirrored by the recovery of the typical spine density after 2 weeks (Fig. 5F) and the restoration of synaptic alterations in the hippocampal CA1 region after 4 weeks: LTP amplitude (Fig. 5G), EPSC amplitudes (Fig. 5H) and AMPA/NMDA ratio (Fig. 5I), with the exception of sEPSC amplitude (Fig. 5L). On the other hand, we report that PLX-induced astrogliosis did not recover during the time window considered, indicating a prolonged alteration of tissue homeostasis following microglia depletion (Fig. 5M).

**Figure 5.**
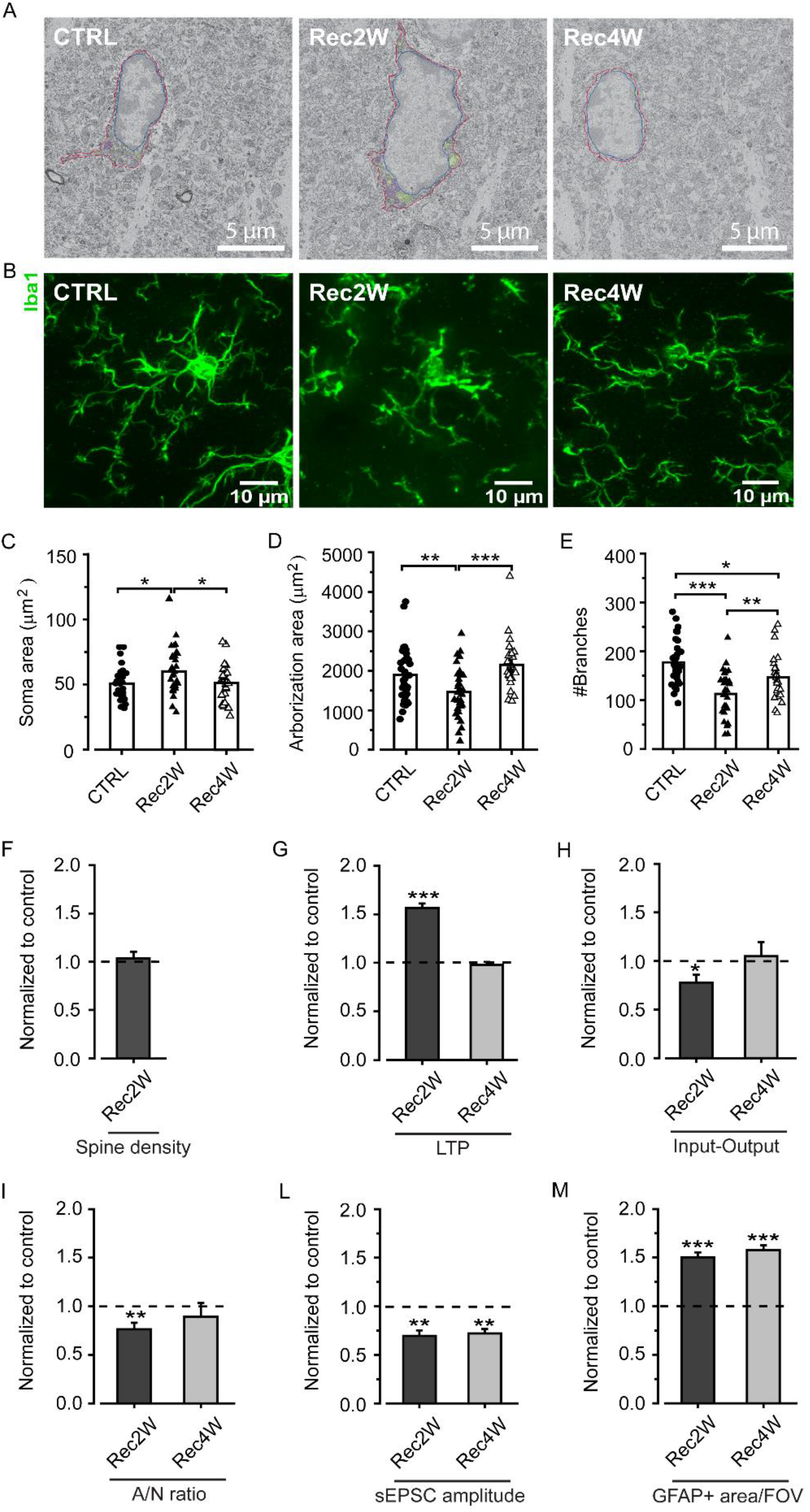
Effects of microglia repopulation on synaptic functions. A) Electron micrograph of microglia in control, 2 and 4 weeks recovery groups in the *stratum radiatum* of the hippocampal CA1. Red line = cytoplasm, blue line = nucleus, purple = mitochondria, green = phagocytic elements. Scale bar = 5 µm. B) Representative confocal images showing morphological features of Iba1-positive microglia in control, 2 and 4 weeks recovery groups in the *stratum radiatum* of the hippocampal CA1. Scale bar = 10 µm. C-E) Quantitative analysis of microglial soma area (C; One-way ANOVA: F(2,90)=4.06, p<.05. Holm-Sidak post-hoc: *p<.05), arborization area (D; One-way ANOVA: F(2,90)=8.1, p<.001. Holm-Sidak post-hoc: **p<.01 ***p<.001) and number of branches (E; One-way ANOVA: F(2,90)=16.06, p<.001. Holm-Sidak post-hoc: *p<.05 **p<.01 ***p<.001) in control (n=34 cells/13 slices/3 mice), 2 weeks recovery (n=34 cells/7 slices/3 mice) and 4 weeks recovery (n=27 cells/7 slices/3 mice) groups. F-I) Analysis of dendritic spine density (F), LTP (G; t-test Rec2W *vs*. ctrl: t=-6.19, ***p<.001), input-output (H; t-test Rec2W *vs*. ctrl: t=-2.74, *p<.05) and AMPA/NMDA ratio (I; t-test Rec2W *vs*. ctrl: t=-3.53, **p<.01) during microglia repopulation. L-M) On the contrary, microglia repopulation did not restore sEPSC amplitude (L; t-test Rec2W *vs*. ctrl: t=2.97, **p<.01; t-test Rec4W *vs*. ctrl: t=-2.98, **p<.01) and astrogliosis (M; t-test Rec2W *vs*. ctrl: t=-6.61, ***p<.001; t-test Rec4W *vs*. ctrl: t=-7.92, ***p<.001) to control condition. Data in F-M) presented as normalized to control values. Data shown as Mean ± SEM.

To ascertain that synaptic recovery was specifically due to microglia repopulation and not merely time-dependent, we measured EPSC amplitudes in conditions of continuous PLX5622 treatment at the same time points chosen for recovery (3 and 5 weeks of treatment). We show that when PLX5622 treatment was not discontinued, recovery of synaptic function was not observed (Fig. EV3).

### Microglial depletion causes a reversible impairment in the acquisition of the NOR task

We tested mice in the NOR task in order to investigate the effects of microglial depletion on learning and memory processes (Fig. 6A). We observed that PLX-treated mice did not habituate to the exploration of the two identical objects across sessions, whereas the control group showed a progressive reduction of object exploration time across trials (Fig. 6B), naturally due to a decrease in novelty after repeated exposure. During the test, PLX-treated mice explored the familiar object as much as the novel one, showing no preference between them, whereas the control group showed a clear preference towards the novel object (Fig. 6C). Consistently, PLX-treated mice showed a significantly lower Discrimination Index (D.I.) compared to control mice (Fig. 6D). To assess whether the deficit in the NOR acquisition could be rescued after microglia repopulation, we switched PLX-treated mice (n=4) to the control diet for 6 weeks and then re-trained them in the NOR task following the same behavioral procedure with a different set of objects. Microglia-repopulated mice recovered the ability to discriminate between the familiar and novel object after re-training, as shown by the higher time spent exploring the novel object (Fig. EV4A) and the higher DI (Fig. EV4B) during the acquisition test session.

**Figure 6.**
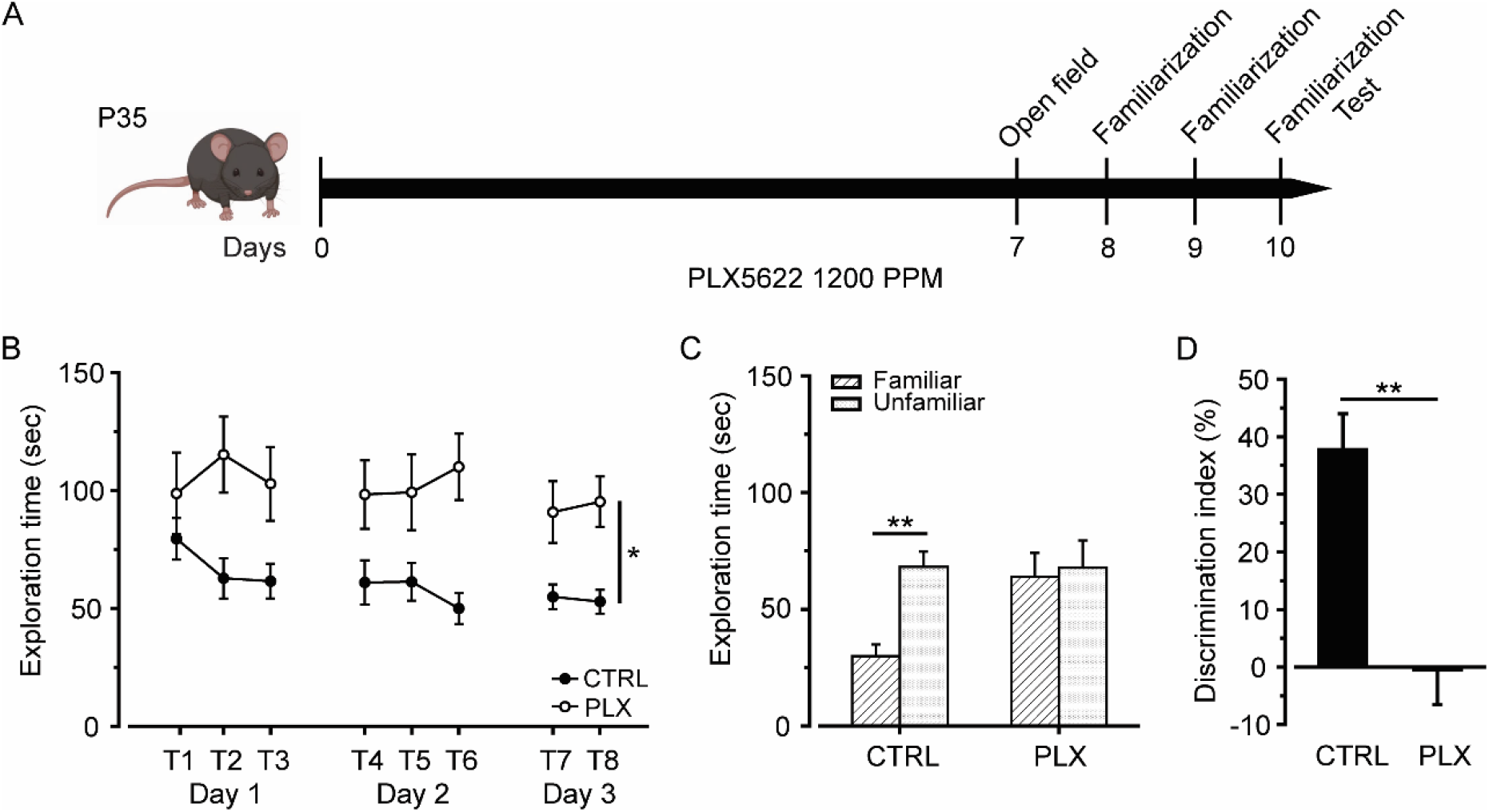
Microglia depleted mice show learning deficits in the NOR task. A) Experimental design for behavioral analysis in (B-D). 35-days-old mice (control n=18, PLX-treated n=11) are fed with PLX5622 chow for 10 consecutive days. After 7 days of treatment, mice are placed in the open field arena to evaluate the locomotor activity. From day 8 to 10, mice are allowed to familiarize with two identical objects during repeated training sessions. On day 10, 1 h after the last training session, mice were tested in the NOR task. B) Exploration time of the two objects across the familiarization sessions. As shown in the graph, PLX-treated mice show increased exploratory behavior compared to control mice. ANOVA for repeated measures, F(1,182) = 11.652; p<.01. C) Bar graph representing the exploration time for the familiar and unfamiliar object during the acquisition test. Two-way ANOVA (group x object), **p<.01. D) Control mice show a higher discrimination index (D.I.) compared to PLX-treated mice, demonstrating to distinguish between novel and familiar object. t-test, **p<.01. Data presented as Mean ± SEM.

We did not find differences between PLX-treated and control mice on either the distance traveled, the velocity or the number of crossings to different areas of the arena (Fig. S4A-D), the time spent grooming nor the time spent in the center and periphery of the open field (Fig S4E-G), which excluded any effect of PLX5622 on general locomotor activity and anxiety-like behavior. Our results demonstrate that microglial depletion impaired the acquisition of the NOR task, pointing to a pivotal role of microglia in hippocampal-dependent cognitive performance.

## DISCUSSION

In this study, we investigated the effects of PLX-induced microglial depletion on hippocampal synaptic function in the adult brain, reporting four main findings. First, in the almost complete absence of microglia, glutamatergic synapses undergo a regression to immature properties, associated with a reduction in functional efficacy and an increase in synaptic plasticity. Second, microglial depletion reduces spine density without affecting spine morphology. Third, microglia-depleted mice fail to acquire the NOR task. Fourth, the observed structural, functional and behavioral deficits are rescued after microglial repopulation. We conclude that microglia are necessary for the maintenance of synaptic function in adulthood.

### Hippocampal CA1 synapses display immature features and enhanced plasticity after microglial depletion

We showed that a short microglial depletion (one-week) causes synaptic alterations in the adult mouse hippocampus. In particular, we observed a reduced efficacy and immature properties in CA1 glutamatergic synapses, as showed by the lower AMPA/NMDA ratio (Hsia *et al*, 1998), reduced synaptic multiplicity (Hsia *et al*, 1998; Paolicelli *et al*, 2011) and higher paired pulse ratio, which is typically associated with a decrease in presynaptic release probability (Dobrunz & Stevens, 1997; Basilico *et al*, 2019). We did not observe alterations in the post-synaptic sensitivity to the neurotransmitter (similar mEPSC amplitude in control and PLX mice), pointing to an alteration at the pre-synaptic level after microglial depletion (Dittman & Regehr, 1996). Altogether, these findings outline a scenario in which, in the absence of microglia, synaptic properties regress, resembling those found in immature glutamatergic CA1 synapses during early postnatal development (Basilico *et al*, 2019). An alternative explanation is that the absence of microglia is associated with a defined pattern of synaptic properties similar to those present during development. Consistently with this hypothesis, the properties of hippocampal CA1 synapses following microglial depletion with PLX5622 administration closely resemble those observed in a dysfunctional microglia model (Basilico *et al*, 2019). In this regard, it should be emphasized that PLX treatment did not produce any significant effect on synaptic properties of mice lacking the fractalkine receptor in microglia (C*x3cr1*^-/-^ mice), indicating an overlapping effect between microglia removal and a model of dysfunctional neuron-microglia signaling (Basilico *et al*, 2019). However, recent evidence reported that microglia may negatively modulate neuronal activity in the striatum (Badimon *et al*, 2020). Indeed, microglial ablation amplifies the activity of neurons upon dopamine D1 receptor agonist treatment, pointing to a potential differential role of microglia in regulating synaptic functioning in a context and brain region-specific manner (Badimon *et al*, 2020).

We also observed a significant increase in LTP amplitude at hippocampal CA1 synapses in microglia-depleted mice. In line with the hypothesis described above, increased LTP was also observed in C*x3cr1*^-/-^ mice (Maggi *et al*, 2011) and it is consistent with the presence of immature synapses. Indeed, a higher level of synaptic plasticity is a general phenomenon in the developing brain (Feldman *et al*, 1999; Hogsden & Dringenberg, 2009), and higher LTP magnitude has been previously reported in developing rodent CA1 (Dumas & Foster, 1995; Dumas, 2012). At the mechanistic level, we may speculate that enhanced plasticity after microglial depletion could rely on the inclusion of ‘immature’ NMDAR subunits at CA1 synapses (i.e. NR2B). Indeed, a decrease of NR2A/NR2B ratio lowers the threshold for LTP (Yashiro & Philpot, 2008), and alterations in the density of NMDAR subunits has been observed after LPS-induced microglial activation (Ma *et al*, 2014).

Although LTP is largely accepted as a mechanism underlying the encoding and storage of memories (Martin *et al*, 2000), a nonlinear relationship between LTP magnitude and learning has been proposed (Barnes *et al*, 1994; Diógenes *et al*, 2011). We can speculate that the elevated LTP amplitude observed in microglia-depleted mice (i.e. dysfunctional LTP or LTP saturation) could contribute to the impairment in the acquisition of the NOR task. In line with this hypothesis, it has been previously demonstrated that inhibiting neuronal-microglial fractalkine signaling, increases LTP amplitude leading to a lack of responsiveness to environmental stimuli (Maggi *et al*, 2011; Milior *et al*, 2016).

### Microglial depletion reduces the number of glutamatergic synapses in the hippocampus

The mechanisms underlying the synaptic dysfunction observed in microglia-depleted mice may partially rely on structural changes. Indeed, we observed a reduction of both spine density and the levels of the post-synaptic marker drebrin in mice treated with PLX. Drebrin A is an actin binding protein distributed in a dot-like pattern along the dendritic protrusions in the murine brain consistent with a dendritic spine localization (Ivanov *et al*, 2009). It is worth noting that drebrin expression is regulated by glutamatergic activity (Takahashi *et al*, 2009). These data along with the decrease in both the AMPA and NMDA components of EPSC is coherent with a reduction in the number of functional glutamatergic synapses in microglia-depleted mice. Similarly to what we observed in the hippocampus, Parkhurst and colleagues showed that microglial depletion reduces learning-dependent formation of postsynaptic dendritic spines in the motor cortex (Parkhurst *et al*, 2013). In contrast, Rice and co-workers reported no change in the number of PSD-95 puncta in hippocampal CA1 neurons after 30 days of PLX3397-induced microglial depletion in control mice and an increase in PSD-95 in microglia-depleted mice through neuronal lesion (Rice *et al*, 2015). In addition to differences possibly arising from the duration of the treatment, this discrepancy is likely to be ascribed to the different impact of microglial removal in basal or inflammatory conditions. Recently Ma and colleagues (Ma *et al*, 2020) also reported an increase in spine density in the developing visual cortex following 2 weeks of PLX3397 treatment. Most likely, this effect is related to the specific developmental window of the treatment, when spine turnover represents a major phenomenon (Cruz-Martin *et al*, 2010). Our results showing a reduction of dendritic spines and a reduced AMPA/NMDA ratio in microglia-depleted mice support the notion that microglia control synaptic functioning and stabilization in the adulthood.

### Recovery of synaptic function following microglial repopulation

To further understand the role of microglia on synaptic functioning, we monitored the functionality of hippocampal synapses after discontinuation of PLX5622 treatment. After two weeks of PLX withdrawal, we observed microglial repopulation of the brain parenchyma, as previously reported (Elmore *et al*, 2014; Zhan *et al*, 2019), and the gradual re-acquisition of their morphological features typical of physiological condition (up to four weeks after PLX removal), following a trend similar to that previously described (Elmore *et al*, 2015; Zhan *et al*, 2019). Remarkably, we observed a functional recovery of synaptic properties in parallel with the recovery of the typical microglial morphology. In particular, two weeks after PLX withdrawal, when microglia re-gained normal density (though with lower branching), the typical spine density was re-established. At this stage, hippocampal CA1 synapses were still weak, immature and more plastic, as demonstrated by the reduced synaptic efficacy, the lower AMPA/NMDA ratio and the higher LTP, all indicative of an ongoing process of maturation. After four weeks of microglial repopulation, we observed a full functional synaptic recovery, with the only exception of the amplitude of the spontaneous EPSC that remained low. Interestingly, the synaptic functional recovery coincided with complete morphological recovery of microglia, but not with the resolution of the astrogliosis, still present at this stage. The recovery of the synaptic function was specifically related to the repopulation and recovery of microglia and not simply an effect of time or age, as aged-matched mice with prolonged PLX administration did not show functional recovery of synapses even after two months.

### Microglial depletion impairs recognition learning in mice

We also report that microglia-depleted mice show a deficit in the acquisition of object recognition memory (NOR), as revealed by the failure in discriminating between the novel and familiar objects during the test. Notably, upon microglia recovery after the discontinuation of the PLX treatment, mice recovered the ability discriminate the novelty. Microglia-depleted mice also displayed enhanced exploration of both objects and did not habituate across training sessions. Consistent with our results, one week of PLX5622 treatment impaired spatial memory acquisition in adult mice (Torres *et al*, 2016). Similarly, one week of microglial depletion, achieved using the tamoxifen-inducible CX3CR1^CreER^ to drive diphtheria toxin receptor expression in microglia, impaired learning in both fear conditioning and the NOR task (Parkhurst *et al*, 2013). Moreover, it has been recently shown that microglial repopulation in aged animals has beneficial effects on cognition, by improving spatial memory and reversing age-dependent deficits in synaptic plasticity (Elmore *et al*, 2018). Overall, our results strongly support the hypothesis that microglia are actively involved in the regulation of learning related processes. It should be noted that the deficit in the NOR task is not explained by alterations in spontaneous locomotor activity or anxiety, consistently with previous findings that PLX treatment does not alter these parameters in different behavioral procedures (Dagher *et al*, 2015; Elmore *et al*, 2014). However, our results and the above mentioned studies are at odds with findings of an intact spatial memory in healthy mice treated with PLX for longer periods (up to 3 weeks) (Dagher *et al*, 2015; Elmore *et al*, 2014) or evidence supporting the PLX-restorative effects on spatial memory deficits in an aged mouse model of Alzheimer’s disease (Dagher *et al*, 2015). Whilst the restorative effects of microglial depletion in animal models of neurological disorders are likely resulting from a decreased inflammation process (Dagher *et al*, 2015; Rice *et al*, 2015), we speculate that in healthy animals the effects of microglial depletion on learning-related processes probably depend on the duration of microglial removal or the cell-specificity of the depletion method. It should be also noted that prolonged inhibition of CSF1R might have systemic effect on hematopoiesis and macrophage functions (Lei *et al*, 2020).

### Microglia are necessary for the maintenance of hippocampal synaptic function and learning in adulthood

In summary, in the absence of normal microglia-neuron interactions, hippocampal CA1 glutamatergic activity is depressed, with weak and less reliable synapses. Functional changes primarily depend on reduced glutamate release and likely determine the reduction of the number of functional synapses and the boost in plasticity mechanisms, possibly associated with a circuit rearrangement, and leading to learning impairment. Indeed, the proper functioning of hippocampal CA3-CA1 synapses is of critical importance for an efficient recognition learning (Broadbent *et al*, 2010; Cohen *et al*, 2013; Cohen & Stackman, 2015). Accordingly, rats with hippocampal lesions display higher rates of object exploration time and lack of habituation during training in the NOR task, similarly to our microglia-depleted mice (Broadbent *et al*, 2010). Our results suggest that the role of microglia on supporting hippocampal excitatory synaptic function is critical for learning and memory. However, our findings should be interpreted with caution because our microglia depletion strategy is systemic and is therefore likely that deficits in other structures might have concurred to the performance in the NOR task. Indeed, beside the hippocampus, other brain structures relevant to the NOR task, such as the perirhinal cortex (Winters *et al*, 2008), might have concurred to the behavioral performance observed.

### Methodological limitations

When discussing our results, it has to be considered that the strategy used to deplete microglia may also have undesired side effects. Indeed, the treatment with CSF1R antagonist is systemic and not restricted to a precise brain area. However, it has been previously shown that PLX5622 effects in the CNS are limited to microglia and do not affect mice general health conditions (Torres *et al*, 2016; Elmore *et al*, 2014). Although prolonged PLX treatments may have effects on peripheral immune cells (Lei *et al*, 2020), PLX treatment do not cause peripheral myeloid cells infiltration in healthy conditions (Rice *et al*, 2015) and repopulation occurs by self-renewal (Zhan *et al*, 2019). We considered the possibility that PLX5622 microglial depletion may cause direct or indirect effects on brain homeostasis due to microglia cell death. Our analyses do not indicate the presence of neuroinflammatory processes in the hippocampus of microglia depleted mice, based on the reduced expression of inflammatory genes. In addition, we did not observe ultrastructural signs of cell death (shrunken size, electron-dense cells) or the appearance of dark microglia, previously associated with ultrastructural signs of oxidative stress (e.g. condensation of the nucleo- and cytoplasm, dilatation of the reticulum endoplasm, mitochondrial alterations) and found in regions where synaptic loss is exacerbated (e.g. in *stratum radiatum* in Alzheimer’s disease and chronic stress (Bisht *et al*, 2016). Furthermore, microglial depletion causes reactive astrogliosis, as indicated by the increase of GFAP signal density and the evidence of high metabolic activity and phagocytosis in astrocytes. We speculate that in the absence of microglia, brain parenchyma homeostasis may require the involvement astrocytes. In particular, given that astrocytes release and regulate several neuroactive molecules that can affect neuronal activity and modulate synaptic plasticity (Perez-Alvarez *et al*, 2014; Ota *et al*, 2013; Allen & Barres, 2005), we cannot exclude a contribution of astrocytes to the alterations in synaptic plasticity observed in the absence of microglia.

In addition, our results should be discussed in light of previous evidence, showing that microglial depletion and repopulation may exacerbate (Rubino *et al*, 2018) or favor the resolution (Han *et al*, 2019) of different neuroinflammatory conditions. Indeed, during microglial repopulation, the brain parenchyma is going through an unnatural developmental process, where microglia although present and responsive (Elmore *et al*, 2015) are gradually re-acquiring their abilities of modulating synapses. Still, the study of synaptic recovery after microglia depletion can be informative about the importance of microglia-neuron interaction in brain plasticity and brain responses after traumatic or inflammatory events.

Our findings clearly demonstrate that microglia are necessary to stabilize the ‘adult’ properties of hippocampal glutamatergic synapses. We provide evidence that the synaptic changes associated with microglial depletion are mechanistically connected to the absence of microglia, as demonstrated by: i) the temporal coincidence of microglial repopulation and morphological recovery with the recovery of synaptic functions and ii) the absence of effect of PLX on synapses functionality in *Cx3cr1*^*-/-*^ mice, iii) the absence of cell death and neuroinflammation.

## Conclusions

Our study highlights the importance of microglia in regulating synaptic functioning in the adult brain. In the absence of microglia, glutamatergic synapses work less efficiently and display immature properties, ultimately affecting cognitive abilities. We conclude that microglia are central in the control of excitatory synaptic activity: besides participating to synaptic development, they are required for the maintenance of glutamatergic synaptic function in adulthood.

## MATERIAL AND METHODS

### Animals

Wild type (WT) C57BL6/J mice were used for the morphological, electrophysiological and behavioral assessments. Thy1::GFP; Cx3cr1::cre ERT2; R26CAG-tdtomato mice were used for dendritic spine analysis. *Cx3cr1*^*-/-*^ mice were used for electrophysiological analysis. All experiments were performed on male mice starting treatments at 5 weeks of age. Mice were maintained under 12-h light/dark cycle (light on at 7 AM) with food and water *ad libitum*. All the procedures followed the guidelines of the national law (DL 26/2014) on the use of animals for research based on the European Communities Council Directive (2010/63/UE), and approved by the ethics committee of the Italian Ministry of Health.

### Microglial depletion

Microglia were pharmacologically depleted by PLX5622, a selective inhibitor of CSF1R, essential for microglia proliferation, differentiation and survival (Elmore *et al*, 2014). PLX5622 was kindly provided by Plexxikon Inc. (Berkeley, USA) and formulated in standard chow at 1200 PPM by Research Diets (refer to the Specific Experiments section for details on diet administration). Drug-free standard chow (Research Diets) was used in control experiments and during PLX5622 withdrawal.

### Experimental overview

#### Experiment 1. Effect of microglial depletion on astrogliosis and neuroinflammation in the hippocampal CA1

In experiment 1 (Exp. 1) we investigated whether the microglial depletion causes an alteration of the brain inflammatory state. We treated three different cohorts of adult C57BL6/J mice (5 weeks old) with either control diet or PLX5622 supplemented chow for 7 consecutive days. One cohort was used for the immunofluorescence analysis of microglial (Iba1+) and astrocytic (GFAP+) markers. The second cohort was used for the ultrastructural analysis of the neuropil, microglia and astrocytes by scanning electron microscopy. The third cohort was used to perform a gene expression profiling of hippocampi with Nanostring analysis.

#### Experiment 2. Effect of microglial depletion on glutamatergic synaptic activity in the hippocampal CA1

In experiment 2 (Exp. 2) we evaluated the excitatory postsynaptic currents recorded from CA1 pyramidal neurons in adult C57BL6/J mice treated with control or PLX diet as described in Exp. 1 (refer to whole-cell patch clamp recordings section for details).

The whole-cell patch clamp recording experiments were also performed in a group *Cx3cr1*^-/-^ mice (mice lacking the microglial fractalkine receptor), treated with control or PLX diet as described in Exp. 1.

#### Experiment 3. Effect of microglial depletion on hippocampal CA1 plasticity

In experiment 3 (Exp. 3) we evaluated synaptic plasticity in the hippocampal CA1 region by inducing LTP in adult C57BL6/J mice treated with control or PLX diet as described in Exp.1 (refer to extracellular field recordings section for details).

#### Experiment 4. Effect of microglial depletion on structural plasticity in the hippocampal CA1

In experiment 4 (Exp. 4) we investigated dendritic spine density/morphology and the distribution of synapses in the hippocampal CA1 of mice treated with control or PLX diet as described in Exp. 1. We took advantage of the Thy1::GFP; Cx3cr1::cre ERT2; R26CAG-tdtomato mouse line (triple transgenic mice expressing green fluorescent protein (GFP) in sparse excitatory neurons (Thy1::EGFP) and tdTomato in microglia (Cx3cr1::CreER16; RC::LSL-tdTomato) (Weinhard *et al*, 2018) for the analysis of the spines (refer to the analysis of spine density and morphology section). For the quantification of pre- and postsynaptic sites we used C57BL6/J (refer to presynaptic and postsynaptic signals section).

#### Experiment 5. Effect of discontinuation of PLX5622 treatment on synaptic function, spine density and microglial morphology in the hippocampal CA1

PLX-induced microglia depletion is reversible after treatment discontinuation (Elmore *et al*, 2014; Zhan *et al*, 2019).

In experiment 5 (Exp. 5) we evaluated synaptic function and plasticity, spine density and microglial morphology 2 and 4 weeks after the discontinuation of PLX treatment. We treated four cohorts of C57BL6/J mice (5 weeks old) as described in Exp. 1, and then switched the mice to a regular diet for two (Rec2W) or four (Rec4W) weeks (Fig. 1). The first cohort was used for the analysis of microglial density and gross morphology by performing immunofluorescence against the Iba1 marker. The second cohort was used for the ultrastructural analysis by scanning electron microscopy. The third cohort was used for extracellular field recordings. The fourth cohort was used for single-cell recordings of excitatory postsynaptic currents in CA1 pyramidal neurons. To this last cohort, we added two extra control groups (mice treated with PLX for 3 and 5 weeks) in order to exclude that the recovery in the synaptic function after PLX5622 withdrawal was due to the mere passage of time (Fig. EV3).

#### Experiment 6. Effect of microglial depletion on the Novel Object Recognition Task

In experiment 6 (Exp. 6) we determined whether the depletion of microglia in healthy adult mice is associated with cognitive deficits. We treated C57BL6/J mice (5 weeks old) with either control diet or PLX5622-supplemented chow for 10 consecutive days. After 7 days of treatment, we initiated the NOR task (Fig. 6). To determine whether PLX induced long-lasting effects, a subset of animals from the PLX-treated group were returned to the control diet over a 6 weeks period, and then re-trained in the Novel Object Recognition Task using the same behavioral procedure with a different set of objects (Fig. EV4).

### Electrophysiology

#### Extracellular field recordings

Acute hippocampal slices were obtained from adult mice (6-10 weeks old). Animals were decapitated under halothane anesthesia (Sigma-Aldrich, Milan, Italy). Brains were rapidly extracted and immersed in ice-cold ACSF solution continuously oxygenated with 95% O_2_ and 5% CO_2_ and containing (in mM): NaCl 125, KCl 4, CaCl2 2.5, MgSO4 1.5, NaH2PO4 1, NaHCO3 26 and glucose 10. Horizontal 350 μm thick slices were cut at +4°C using a vibratome (Thermo Scientific, USA). Slices were transferred into an incubation chamber containing oxygenated ACSF and allowed to recover for 1 h at 30 °C. Recordings were performed at 30-32 °C under constant perfusion of oxygenated ACSF at 2ml/min rate within a time window of 1–6 h after slice preparation. We placed a concentric bipolar stimulating electrode (SNE-100 × 50 mm long, Elektronik–Harvard Apparatus GmbH) in the hippocampal *stratum radiatum* to stimulate Schaffer collateral fibers. Stimuli consisted of 100 μs constant current pulses of variable intensity, applied at 0.05 Hz. The recording glass micropipette (0.5–1 MΩ) was filled with ACSF and placed in the CA1 hippocampal region, at 200–600 μm from the stimulating electrode, in order to measure orthodromically-evoked field extracellular postsynaptic potentials (fEPSP). The stimulus intensity was adjusted to evoke fEPSPs of amplitude about 50% of maximal amplitude with minimal contamination by a population spike. We monitored evoked responses online, and stable baseline responses were recorded for at least 10 min. Only slices that showed stable fEPSP amplitudes were included in the analysis. LTP was induced by high-frequency stimulation (HFS, 1 train of stimuli at 100 Hz, of 1 s duration). To analyze the time course of fEPSP amplitude, the recorded fEPSP was routinely averaged over 1 min (n=3 traces). We calculated fEPSP amplitude changes following the LTP induction protocol with respect to the baseline (30 min after *vs* 1 min before LTP induction). fEPSPs were recorded and filtered (low pass at 1 kHz) with an Axopatch 200A amplifier (Molecular Devices, LLC) and digitized at 10 kHz with an A/D converter (Digidata 1322A, Axon Instruments). Data were analyzed off-line with Clampfit 10 software (Molecular Devices, LLC).

#### Whole-cell patch clamp recordings

Acute hippocampal slices were obtained from adult mice (6-10 weeks old). Animals were decapitated under halothane anesthesia (Sigma-Aldrich, Milan, Italy). Brains were rapidly extracted and immersed in ice-cold ACSF, containing (in mM): KCl 2.5, CaCl_2_ 2.4, MgCl_2_ 1.2, NaHPO_4_ 1.2, glucose 11, NaHCO_3_ 26, glycerol 250, continuously oxygenated with 95% O_2_ and 5% CO_2_ to maintain the physiological pH. Horizontal 250 μm thick slices were cut at 4°C, using a vibratome (DSK, Dosaka EM, Kyoto, Japan) and placed in a chamber filled with oxygenated ACSF containing (in mM): NaCl 125, KCl 2.5, CaCl_2_ 2, MgCl_2_ 1, NaHPO_4_ 1.2, NaHCO_3_ 26 and glucose 10. Before use, slices were allowed to recover for at least 1 h at room temperature (24 ± 1 °C). All recordings were performed on slices submerged in ACSF and perfused with the same solution in the recording chamber. In order to increase the release probability, where indicated, the extracellular Ca^2+^/Mg^2+^ concentration ratio of ACSF was increased to 8 (Ca^2+^ = 4 mM, Mg^2+^ = 0.5 mM). We recorded spontaneous and evoked excitatory postsynaptic currents (EPSCs) from hippocampal CA1 pyramidal neurons at -70 mV using a patch clamp amplifier (Axopatch 200A, Molecular Devices, LLC). Data were filtered at 2 kHz, digitized (10 kHz), acquired using pClamp 10.0 software (Molecular Devices) and analyzed off-line using Clampfit 10 (Molecular Devices, LLC). Patch pipettes (3-5 MΩ) were filled with intracellular solution containing (in mM): Cs-methanesulfonate 135, HEPES 10, MgATP 2, NaGTP 0.3, CaCl_2_ 0.4, MgCl_2_ 2, QX-314 2, BAPTA 5 (pH adjusted to 7.3 with CsOH). Bicuculline methochloride (10 μM) was added to the extracellular solution to block GABA_A_ receptors. For experiments comparing sEPCSs and mEPSCs, EPSCs were recorded from the same cell (one cell per slice). mEPSCs were recorded after 10 minutes of bath perfusion with tetrodotoxin (TTX, 1 µM). In experiments on evoked synaptic currents, electrical stimulation was applied by theta glass tubes (tip 15-20 µm) filled with external solution. The stimulating electrode, connected to a stimulus isolation unit (Iso-stim A320, WPI), was placed in the hippocampal CA1 *stratum radiatum* to activate the Schaffer collaterals projecting from CA3 to CA1 as previously described (Basilico *et al*, 2019). Synaptic responses were evoked by stimulating for 100 µs at 0.1 Hz; the stimulus intensity was adjusted accordingly to the experiment.

AMPA-mediated EPSCs were evoked by paired pulse stimulations (interval 50 ms), to determine the paired pulse ratio (PPR). For input/output curves, Schaffer collaterals were stimulated at increasing intensities (0.1-10 mA). Each pulse of a given intensity was repeated 6 times, to obtain an average response. The input/output of the AMPA component was determined at -70 mV, whereas the input/output of the NMDA component was determined at +40 mV in presence of NBQX 10 µM. To determine the AMPA/NMDA ratio, stimulus strength was adjusted to obtain at -70 mV stable AMPA-mediated EPSC with an amplitude corresponding to the 50% of the maximum response. NMDA currents were recorded from the same neuron and using the same stimulus strength. For NMDA EPSC, we measured the EPSC reversal potential (ERev) in each cell and corrected the NMDA recording potential consequently (+40 mV ERev), to avoid potential voltage clamp errors. NMDA peak amplitude was measured with a delay of 25 ms from the AMPA peak. The AMPA/NMDA ratio was then calculated using the equation: ratio = AMPA EPSC amplitude/NMDA EPSC amplitude.

For current clamp experiments and sIPSC recordings, 350 µm-thick hippocampal slices were prepared as described above, transferred to a submerged chamber and perfused with oxygenated ACSF (3-4 ml/min; same composition as for sEPSCs). CA1 or CA3 neurons were visualized using the IR-DIC optics of an upright Leica DMLSF microscope and current- or voltage clamp recordings were obtained using a MultiClamp 700B amplifier and the pClamp9 suite (Molecular Devices). All recordings were performed at room temperature (24 ± 1 °C). Patch pipettes (4-10 MΩ) were filled with intracellular solution containing (in mM): 110 K-gluconate, KCl 12, Na2-phosphocreatine 10, HEPES 10, Mg2-ATP 4, Na2-GTP 0.2, EGTA-K 0.1 (pH 7.3 with KOH; osmolarity corrected with sucrose when needed to reach ∼290-310 mOsm); for sIPSC recordings, the solution was the same except for substituting 110 mM K-gluconate with 120 mM CsMetSO3; QX-314 (5 mM) was added daily to avoid AP firing at depolarized potentials.

In current clamp experiments, the sub- and supra-threshold properties of pyramidal CA3 neurons were studied within the first 3 minutes after patch rupture and while holding the membrane potential at approximately -80 mV by constant current injection. Current steps (I_inj_) ranging in amplitude from 200 to 600 pA (50 pA increments; 1 s duration) were injected to estimate the input resistance (Rin) and to build the *f – I* relationship. The rheobase, the amplitude of the injected current necessary to induce the first AP, was estimated graphically (Clampfit) from the voltage response to a series of 5 pA-incremented consecutive steps of I_inj_ (50 ms duration; range: 0 ÷ 0.4 nA).

In voltage clamp, sIPSCs were recorded holding the membrane potential at + 10 mV; estimated reversal potential (Erev) for glutamatergic synaptic currents was 0 mV, thus no blockers were necessary to isolate GABAergic currents (estimated Erev approx. -65 mV). To improve whole-cell voltage clamp, series resistance (Rs) was compensated (range 30-70%) and recordings bearing excessive increase of Rs were excluded from analysis. In some voltage clamp experiments, 10 µM SR95531 (a blocker for GABA_A_ receptors; Tocris) was used to verify the GABAergic nature of the currents recorded (data not shown). Current clamp recordings were filtered online at 10 kHz and sampled at 50 kHz; synaptic currents were filtered at 2 kHz and digitized at 10 kHz (amplifier in-built 8-pole low-pass Bessel filter and Molecular Devices Digidata 1322A, respectively). Recordings used for analysis were post-hoc filtered at 1 kHz.

### Analysis of spine density and morphology

#### Dendritic spines imaging

Mice were transcardially perfused with phosphate buffer 0.9% saline solution (PBS) and 4% paraformaldehyde (PFA) in 0.1 M pH 7.4 phosphate buffer (PB), under halothane anesthesia (Sigma-Aldrich, Milan, Italy). Brains were rapidly removed, post-fixed overnight in PFA 4% and incubated in PBS 30% sucrose solution for 24h at +4°C. Before being frozen at -80°C, brains were placed for at least 2 h in chilled isopentane. We cut the frozen brains into 40 µm horizontal slices using a cryostat (Leica). Once fluorescent neurons of hippocampal CA1 were identified, images of primary dendrites that ramify in secondary and tertiary dendrites in the CA1 *stratum radiatum* were acquired using a confocal laser scanning microscope (iX83 FV1200) with a 60x oil immersion objective. Fluorescence was separated by a dichroic mirror with 572–700 nm emission filter (green channel: for EGFP fluorescence detection) and detected Z-stack images by photomultipliers (1600×1600 pixels, 0.044 µm per pixel, 0.1 µm z-step).

#### Dendritic spines quantification and analysis

To increase resolution, images were deconvoluted (50 iterations, 0.01 of quality change) with the software Huygens Professional 18.04. Analysis was performed blinded to the experimental conditions using the NeuronStudio software. We isolated segments of known length from secondary or tertiary dendrites, and then reconstructed the segment with the software. Dendritic spines were automatically counted and classified into three morphological classes (thin, stubby and mushroom) (Rodriguez *et al*, 2008). Not detected spines were added manually and classified based on shape features as follows: spines displaying a wide protuberance in absence of a neck were classified as stubby, thin and elongated spines with head and neck of similar diameter were classified as thin spines, and spines with with a prominent head and a thin neck were classified as mushroom (Peters & Kaiserman-Abramof, 1970; Hering & Sheng, 2001). To calculate dendritic spine density, spine number was divided by the length of the dendritic segment and expressed as number per micrometer.

### Analysis of presynaptic and postsynaptic signals

#### Immunofluorescence

Under a sub-lethal dose of carbon dioxide, mice were transcardially perfused with cold PBS and 4% PFA in 0.1M PB. Brains were rapidly removed, post-fixed overnight in PFA 4%, washed with PB and cryoprotected in PB 30% sucrose solution. We then collected 40 µm-thick coronal sections with a cryostat microtome (Leica Microsystems) at -20°C. Sections containing the hippocampus were rinsed three times in PB and incubated with blocking solution, containing 10% Bovine serum albumin and 0.1% Normal donkey serum (Sigma Aldrich, St Louis, USA) in PB 0.3% Triton X-100 (Applichem, BioChemica, Darmstadt, Germany) 0.1 M, for 1h at room temperature. Slices were then incubated with primary antibodies in blocking solution overnight at room temperature. We used as presynaptic and postsynaptic markers, respectively: Rabbit anti-Synapsin I polyclonal antibody (1:100, Sigma Aldrich, St Louis, USA) and guinea pig anti-Drebrin polyclonal antibody (1:100, OriGene Technologies, Herford, Germany). Neural dendrites were detected with the neurofilament marker anti-SMI 311 mouse monoclonal antibody (1:600, Millipore Bioscience Research Reagents, USA). The day after, free floating slices were washed (3×10-min) in PB at room temperature and incubated with secondary antibodies for 2h, followed again by three 10-min rinses in PB. We applied DAPI for 5 min during the second rinse, dissolved 1:2000 in PB solution. Control background experiments were performed by incubating secondary antibodies alone.

#### Structured illumination (SIM) microscopy

Image acquisition was performed using a Nikon Eclipse Ti equipped with: X-Light V2 spinning disk combined with a VCS (Video Confocal Super-resolution) module (CrestOptics) based on structured illumination; LDI laser source (89 North); Prime BSI Scientific CMOS (sCMOS) camera with 6.5 µm pixels (Photometrics). The images were acquired with Metamorph software version 7.10.2. (Molecular Devices) with a 100x PlanApo Lambda oil objective (1.45 numerical aperture) and a z-step size of 0.1 µm to obtain a total Z-stack of about 1.2 µm. In order to achieve super-resolution, we processed the raw data obtained by the VCS module with a modified version of the joint Richardson-Lucy (jRL) algorithm (Ingaramo *et al*, 2014; Ströhl & Kaminski, 2015; Chakrova *et al*, 2016), where the out of focus contribution of the signal was explicitly added in the image formation model used the jRL algorithm and evaluated as a pixel-wise linear “scaled subtraction” (Heintzmann & Benedetti, 2006) of the raw signal. *Analysis and quantification*. Presynaptic (Synapsin I marker) and postsynaptic (Drebrin marker) signals were analyzed using the Synapse analyzer plugin of Fiji software (Dzyubenko *et al*, 2016) with the following settings: input dimensionality 3D, triangle threshold, presynaptic and postsynaptic particle size set at 1 px square. We quantified Synapsin+ and Drebrin+ puncta per area.

### Analysis of astrogliosis, microglial density and morphology

#### Immunofluorescence

Mice were perfused and brains sliced as previously described (refer to the analysis of spine density and morphology section). Sections were then rinsed in PBS (3×5 min) and we performed antigen retrieval by soaking slices for 40 min in a warm solution containing (in mM): 10 Na-citrate, 0.05% Tween 20, pH 6.0, 90°C. Subsequently, slices were incubated for 45 min in blocking solution (3% goat serum and 0.3% Triton X-100 in PBS). Primary antibodies diluted in blocking solution were incubated overnight at 4°C (anti-GFAP z0334 DAKO 1:400, anti-Iba1 019-19741 Wako 1:400). After three washes in PBS, we incubated the sections for 45 min with secondary antibodies (Goat anti Rabbit -Alexa Fluor 647, ThermoFisher) and Hoechst in order to visualize nuclei. We then coverslipped them with fluorescence mounting medium S3023 (Dako) and acquired the images by confocal microscopy (See below).

#### Structured illumination (SIM) microscopy

As for the analysis of presynaptic and postsynaptic signals, image acquisition was performed through a Nikon Eclipse Ti equipped with a X-Light V2 spinning disk combined with a VCS (Video Confocal Super resolution) module (CrestOptics) based on structured illumination and with a LDI laser source (89 North). The images were acquired by using Metamorph software version 7.10.2. (Molecular Devices) with a 10x and 60x PlanApo l oil objective (1.4 numerical aperture) and sectioning the slice in Z with a step size of 0.2 µm for spinning disk to obtain a total Z-stack of about 10 µm (for astrocytes measurement) or 25-35 µm for microglia morphology analysis.

#### Astrogliosis analysis

Maximal intensity projection images (50 z-stack planes) were analyzed using Metamorph software. After thresholding the images, astrogliosis was quantified as the area occupied by fluorescent GFAP+ cells versus total field of view.

#### Microglial morphology and density

Image processing was performed using ImageJ software (NIH). To quantify microglial density, the number of Iba1+ cells was reported as number of somas per acquired volume. Maximal intensity projections of Iba1 confocal images were analyzed to obtain morphological indicators of cell complexity, as previously described (Basilico *et al*, 2019). Only cells whose body and processes were entirely contained in the image were included in the analysis. For each cell, the soma area and the arborization domain were assessed. Through skeleton analysis, we calculated the total number of microglial processes.

#### NanoString gene expression

Neuroinflammation mouse panel, containing 757 mouse neuroinflammatory genes and 13 internal reference controls, was used for control and PLX hippocampus samples. Briefly, a total of 100 ng RNA in a volume of 5 μl was hybridized to the capture and reporter probe sets at 65°C for 20 h according to the manufacturer’s instructions. Data were collected using the nCounter Digital Analyzer (NanoString). RNA counts were normalized using the geometric mean of the housekeeping genes included in the panel, after validation against positive and negative controls, using the nSolver 4.0 software (NanoString). Fold changes were calculated comparing untreated (CTRL) and treated (PLX) samples.

### Ultrastructural analysis of the neuropil, microglia and astrocytes

#### Immunohistochemistry for electron microscopy

Four mice per experimental group were anesthetized with sodium pentobarbital (80 mg/kg, i.p.) and perfused with 0.1% glutaraldehyde in PFA 4%. Coronal sections containing the *stratum radiatum* of the hippocampal CA1 were selected based on the stereotaxic atlas Paxinos and Franklin 4th edition. The sections were washed in PBS 50 mM, (pH 7.40) to remove the cryoprotectant solution, incubated 5 min in 0.3% hydrogen peroxide and 30 min in 0.1% sodium borohydride, followed by 1 h long incubation in a blocking buffer solution (10% fetal bovine serum, 3% bovine serum albumin, 0.01% triton X-100). The tissues were then incubated overnight with the primary antibody rabbit anti-Iba1 (1/1000; Wako, catalogue number 019-19741) in the blocking buffer solution at 4°C. The following day, sections were thoroughly washed before being incubated 90 min with the biotinylated secondary antibody goat-anti rabbit (1:300; Jackson ImmunoResearch, catalogue number 111-066-046) at room temperature. Hippocampal sections were incubated 1 h in an avidin-biotin complex solution (1:100, Vector Laboratories) and developed in a tris-buffer solution (TB, 0.05 M, pH 8.0) containing 0.05% 3,3’Diaminobenzidine (DAB) with 0.015% hydrogen peroxide for 5 min. The sections were then processed for scanning electron microscopy. Briefly, sections were incubated 1 h in a solution of equal volume containing 4% osmium tetroxide and 3% potassium ferrocyanide in phosphate buffer (PB, 0.1M, pH 7.4), 20 min in a heated solution of distilled water combined with thiocarbohydrazide and 30 min in an aqueous solution of 2% osmium tetroxide. The sections were then dehydrated with an increasing amount of ethanol (5 min each dehydrating step): 2×35%, 1×50%, 1×70%, 1×80%, 1×90%, 3×100%. Following the dehydration, the tissues were washed in propylene oxide to remove the excess ethanol and were immersed overnight in Durcupan resin. The next day, sections were placed between two ACLAR sheets (Electron Microscopy Sciences) and were left to polymerize for 3 days at 55°C.

#### Scanning electron microscopy (SEM) using array tomography

The hippocampal CA1 region was then dissected from the processed biological samples using a knife and a binocular microscope. Tissues were then mounted on resin blocks using superglue and 70 nm thick ultrathin sections were acquired using a Leica UC7 ultramicrotome equipped with a diamond knife (Diatome). Floating sections were loaded onto dust-free silicones chips (EMS) which were installed on round metal stub (EMS) inserted into a Zeiss Crossbeam 540 focused ion beam scanning electron microscope (FIB-SEM). Once in the FIB-SEM, the hippocampal *stratum radiatum* was identified using backscattered electrons (ESB) and secondary electrons (SE2) detectors at 10 kV for initial focus, then at 1.4kV when samples were at a sufficient working distance to perform high-resolution focus on the brain ultrastructure. These steps were controlled using SmartSEM software (Fibics). Automated imaging of whole sections was performed using ATLAS Engine 5 software (Fibics) and imaging regions of interest were drawn in order to produce 25 nm and 5 nm resolution mosaics for regional analysis and microglial ultrastructural analysis, respectively. Resulting mosaics were then stitched together using Atlas 5 and individually verified for precise title matching. The resulted images were exported as TIFF files using Atlas 5 as well and experimental conditions were blinded to the investigator using ImageJ.

#### Ultrastructural analyses of microglia

Four-to-nine microglial cell bodies were analyzed qualitatively in one animal per condition. For the qualitative analysis of microglial ultrastructure, the following parameters were evaluated: the presence of phagocytic inclusions, elongated or altered mitochondria, lysosomes, lipid bodies, lipofuscin, synaptic contacts, extracellular space and/or digestion and the dilation of the endoplasmic reticulum (ER) and Golgi apparatus. Microglial cell bodies were identified by their immunoreactivity to Iba1, their overall shape, unique heterochromatin pattern, extracellular space pockets as well as their long and narrow ER (Peters *et al*, 1991; Tremblay *et al*, 2010). The analysis of these parameters was based on the transmission electron microscopy analysis of microglia previously published (Bisht *et al*, 2016; Hui *et al*, 2018; El Hajj *et al*, 2019; Savage *et al*, 2019; St-Pierre *et al*, 2019). Briefly, we considered mitochondria elongated if their length was greater than 1000 nanometer and, altered if they appeared swollen, or if broken cristae were observed (Bisht *et al*, 2016; Hui *et al*, 2018). We identified lysosomes by the presence of granules, the association with vesicles or lipid bodies and their electron-dense heterogeneous content (El Hajj *et al*, 2019). We distinguished lipofuscin from lysosomes by their distinct fingerprint-like pattern (Peters *et al*, 1991). We identified lipid bodies by their round shape and homogenous content (Hui et al., 2018). We determined extracellular space and digestion by the presence of electron-lucent content found between the microglia and the parenchyma (Tremblay *et al*, 2012). We characterized the synaptic contact by an interaction with either a pre-synaptic axon terminal and/or post-synaptic dendritic spine recognized by the presence of a post-synaptic density. We determined the dilation of the ER and Golgi apparatus by the electron-lucent pockets found inside the organelles (Hui *et al*, 2018; St-Pierre *et al*, 2019).

### Novel object recognition task

#### Equipment

NOR task was carried out in an open field arena (40 x 40 x 40 cm) made in plexiglass placed in a dimly (30-40 lx) illuminated soundproof room. Stimulus objects were made of ceramic or plastic and varied in color, size and shape. The role (familiar or novel) and the relative position of the objects was counterbalanced and randomly permuted for each mouse. After each session, before starting the new trial, the open field arena and the stimulus objects were cleaned with a 70% ethanol solution to ensure the absence of olfactory cues. Mouse behavior was recorded by a video tracking and analysis system (ANY-maze video-tracking software;5.33; Stoelting Co., Wood Dale, Il).

#### Behavioral Procedure

The experiments were performed blinded to the treatment condition. Mice were handled for 7 consecutive days (1 min/ twice a day) prior to the behavioral assessment to ensure an adequate habituation to the experimenter. On the first day of the experiment, mice were habituated to the open field arena by allowing them to freely explore for 10 min in the absence of any other behaviorally relevant stimulus (twice/ 60 min apart). On the second day, familiarization/training started: mice were placed in the open field arena containing two identical objects and they were free to explore for 10 min (3 trials / 60 min apart). The same procedure was repeated on the third day and during the first two trials on the fourth day. On the fourth day, 60 min after the last familiarization/training trial, mice were subjected to an acquisition test (5 min). For this purpose, one of the objects used during training was randomly replaced by a novel one. Exploration of the object was defined as active exploration with the nose pointing towards the object at a distance smaller than 1 cm. The discrimination index (D.I.) was calculated as the ratio of the difference between the exploration time of the unfamiliar (UF) minus the familiar (F) object and the total exploration time: DI = (UF – F)/Total exploration time * 100 (Broadbent *et al*, 2004). We also recorded the number of crossings, the total distance traveled (meters), the distance traveled in the center of the arena (meters), the average speed (meters/sec) and the time spent grooming.

### Statistical analysis

All data are reported as mean ± SEM. Origin 6 and SigmaPlot 12 (Systat Software Inc., San Jose, CA) softwares were used for statistical analysis. For electrophysiological recordings, n/N refers to numbers of slices/mice (field recordings) and cells/mice (whole-cell patch clamp). Electrophysiological data were analyzed by unpaired t-test, when comparing only two groups, or one-way ANOVA, when comparing more than two groups. Kolmogorov-Smirnov test was used to compare cumulative probability curves. Two-way ANOVA (treatment x stimulus intensity) was used to analyze the input-output curves. Post-hoc comparisons were performed using the Holm–Sidak test. Dendritic spine density and morphology, microglial density and morphology, and astrogliosis were analyzed by one-way ANOVA, followed by Holm–Sidak post-hoc analysis. For Nanostring analysis, we determined significant Differential Expressed (DE) genes by t-test using MeV software considering fold changes higher than 1.5 fold, lower than 0.5 fold and p ≤ 0.05. We performed unsupervised hierarchical clustering and heat map analysis using MeV software. Repeated measure analysis of variance (ANOVA) was used to analyze object exploration time during the familiarization/training trials in the NOR task. Post-hoc comparisons between trials were performed using the Tukey’s test. One-sample t-test was used to verify the control group correctly acquired the task (D.I. different from zero). Unpaired t-test was used to compare locomotor activity and performance in the acquisition test between control and microglia-depleted mice. Normality tests were performed with Origin 6 and non-parametric tests were used when appropriate.

## ACKNOWLEDGEMENTS

The work was supported by a grant from MIUR (PRIN 2017HPTFFC_003) to DR and in part by funds to SDA (CrestOptics-IIT JointLab for Advanced Microscopy) and DC. BB and LF were supported by the PhD program in Clinical-Experimental Neuroscience and Psychiatry, Sapienza University, Rome. The authors thank Alessandro Felici Claudia Valeri and Arsenio Armagno for animal husbandry and colonies maintenance. They also wish to thank Drs Piotr Bregestovski and Michal Schwartz for helpful discussion and criticism. PLX5622 was provided under Materials Transfer Agreement by Plexxikon Inc. (Berkeley, CA, USA).

## AUTHOR CONTRIBUTIONS

BB and LF carried out and analyzed all evoked EPSC and sEPSC experiments. PR performed behavioral experiments with help from IR. LM and MTG performed LTP experiments. MR performed GABAergic and excitability recordings. MR, AG, VF, IR, MCM, SM, LF, MG, VDT, SG, MR and ES performed immunohistochemistry, confocal acquisition, and morphology analysis. MET, MC and MKStP performed EM acquisition and analysis. DS performed nanostring analysis. BB, DR and SDA designed the experiments with input from LF. DR, BB and SDA wrote the article with the help of IR, DC, PR, CTG and CL. DR conceived the project.

## CONFLICT OF INTEREST

The authors declare that they have no conflict of interests

## EXPANDED VIEW FIGURE LEGENDS

**Figure EV1.**
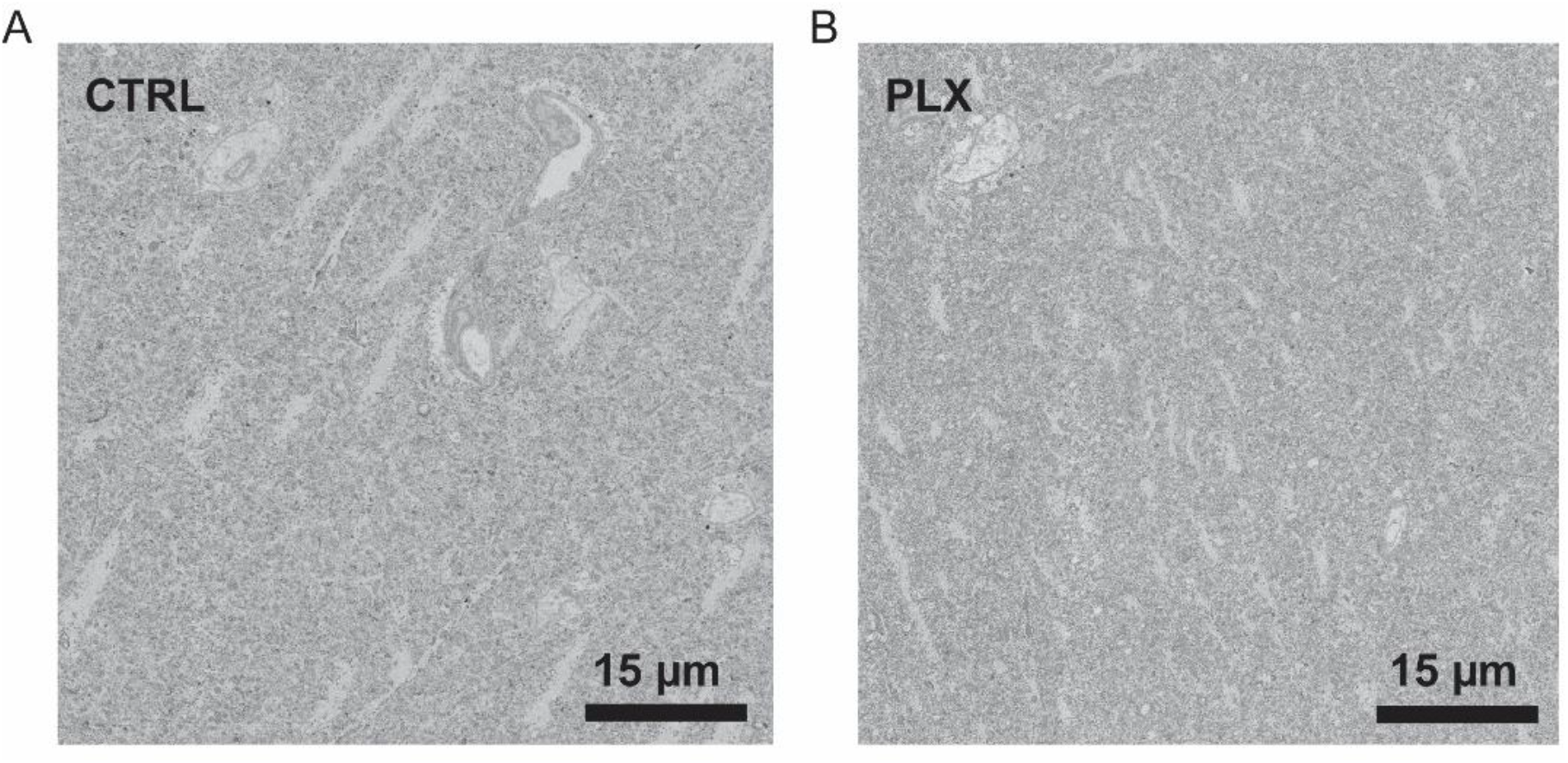
Microglia depletion does not cause tissue damage. A-B) Electron micrograph of the *stratum radiatum* neuropil from a control (A) and PLX-treated (B) mouse showing healthy dendrites alongside astrocytic cells and no ultrastructural signs of neuroinflammation. Scale bars = 15 µm.

**Figure EV2.**
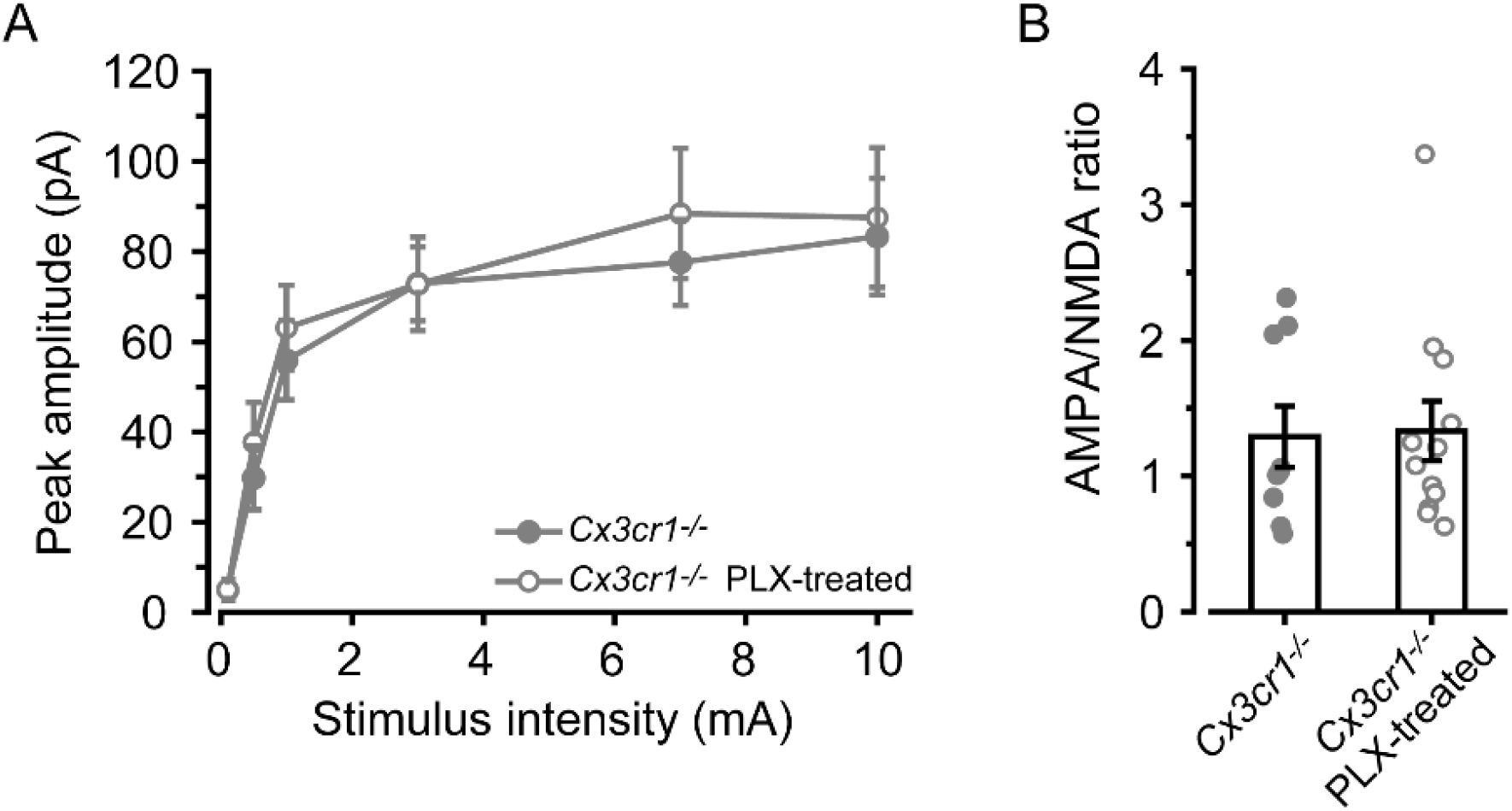
PLX treatment does not affect synaptic transmission in *Cx3cr1*^*-/-*^ mice. A) Input-output curve of evoked EPSC peak amplitudes recorded at −70 mV from *Cx3cr1*^*-/-*^ (n=17 cells/5 mice) and *Cx3cr1*^*-/-*^*-*PLX neurons (n=23 cells/7 mice). No differences detected between the two groups. B) Bar graph showing no differences in AMPA/NMDA ratio from *Cx3cr1*^*-/-*^ (n=9 cells/4 mice) and *Cx3cr1*^*-/-*^ PLX neurons (n=12 cells/5 mice). Data shown as Mean ± SEM.

**Figure EV3.**
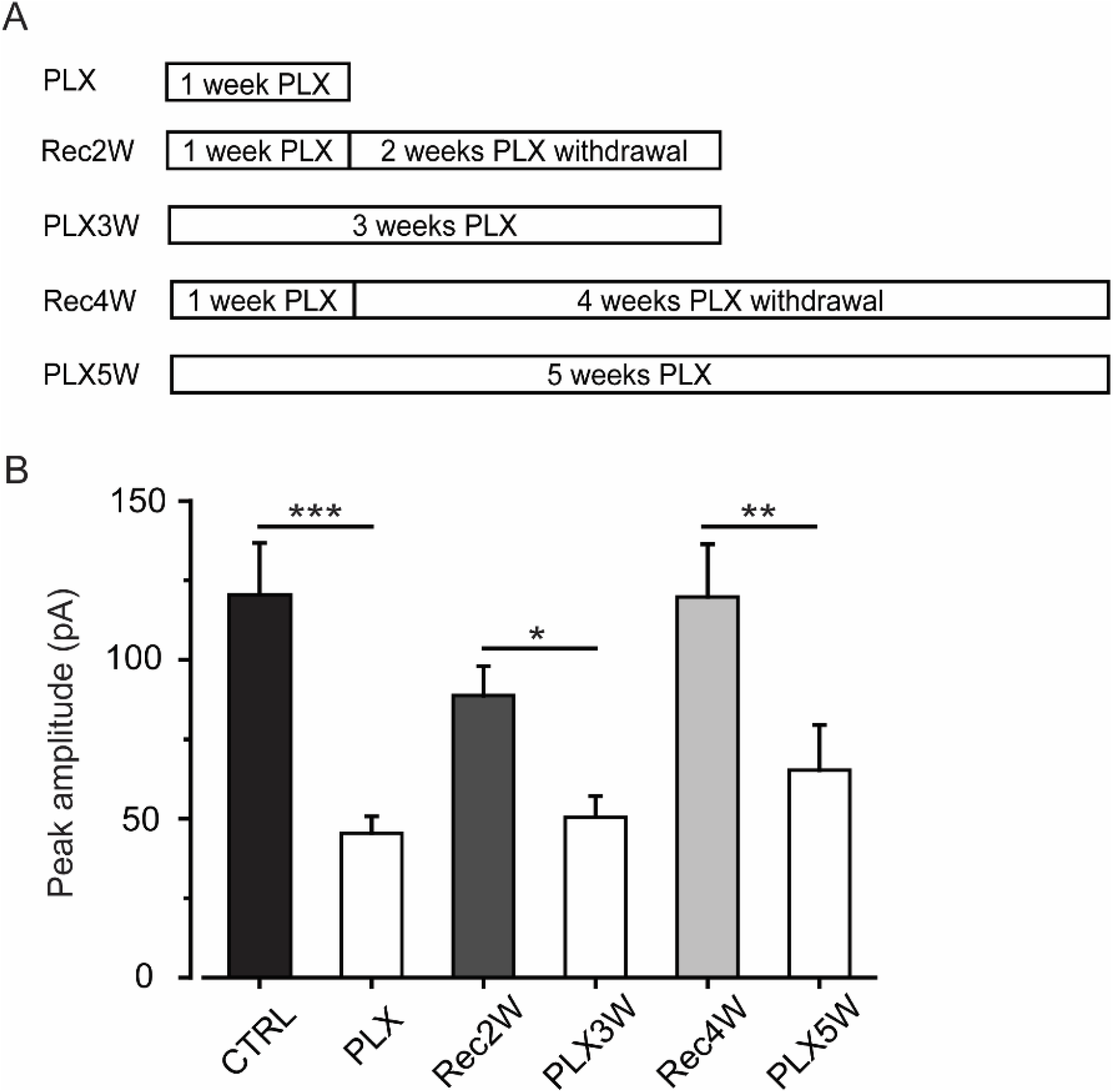
Effect of prolonged microglial depletion on evoked AMPA-mediated currents. A) To assess the potential effect of prolonged PLX treatment on synaptic transmission, additional experimental groups of mice are treated with PLX for 3 (PLX3W) or 5 (PLX5W) weeks. B) Bar graph representing the peak amplitudes evoked by a 7 mA stimulation of Schaffer collaterals and recorded from hippocampal CA1 pyramidal neurons. EPSCs have been extracted from the input/output curve protocol. Control (n=24 cells/8 mice), PLX (n=24 cells/7 mice), Rec2W (n=19 cells/7 mice), PLX3W (n=13 cells/3 mice), Rec4W (n=13 cells/7 mice), PLX5W (n=11 cells/3 mice). One-way ANOVA: F(2,47)=1.5, p<.001; Holm-Sidak post-hoc: *p<.05, **p<.01, ***p<.001. Data presented as Mean ± SEM.

**Figure EV4.**
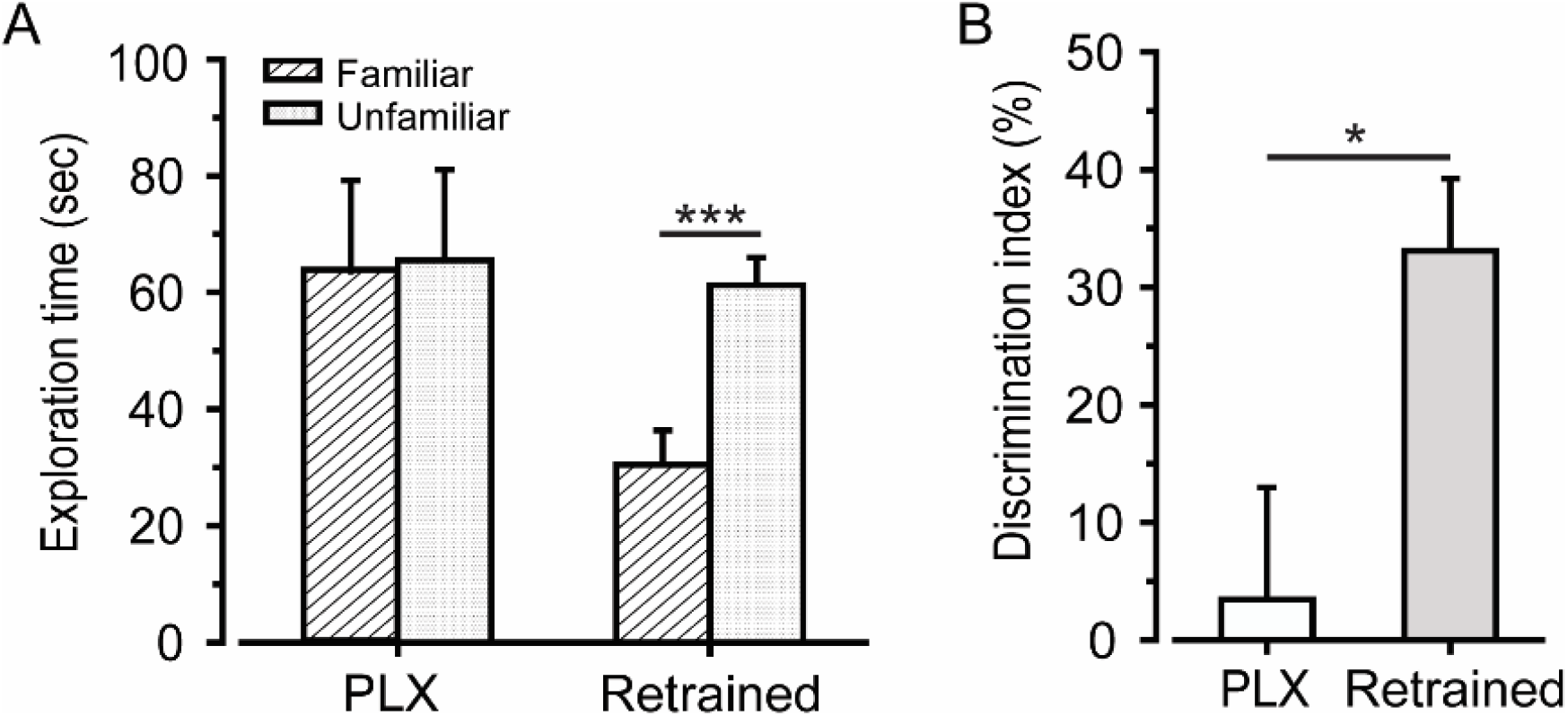
Microglia repopulation rescues the memory deficit observed in the NOR task. A) Bar graph showing the exploration time for the familiar and unfamiliar object during the acquisition test. Two-way ANOVA, ***p<.001. B) Retrained mice (n=4) show a higher discrimination index (D.I.) compared to the first training, demonstrating to distinguish between novel and familiar object. t-test, *p<.05. Data presented as Mean ± SEM.

## APPENDIX FIGURE LEGENDS

**Appendix Figure S1.**
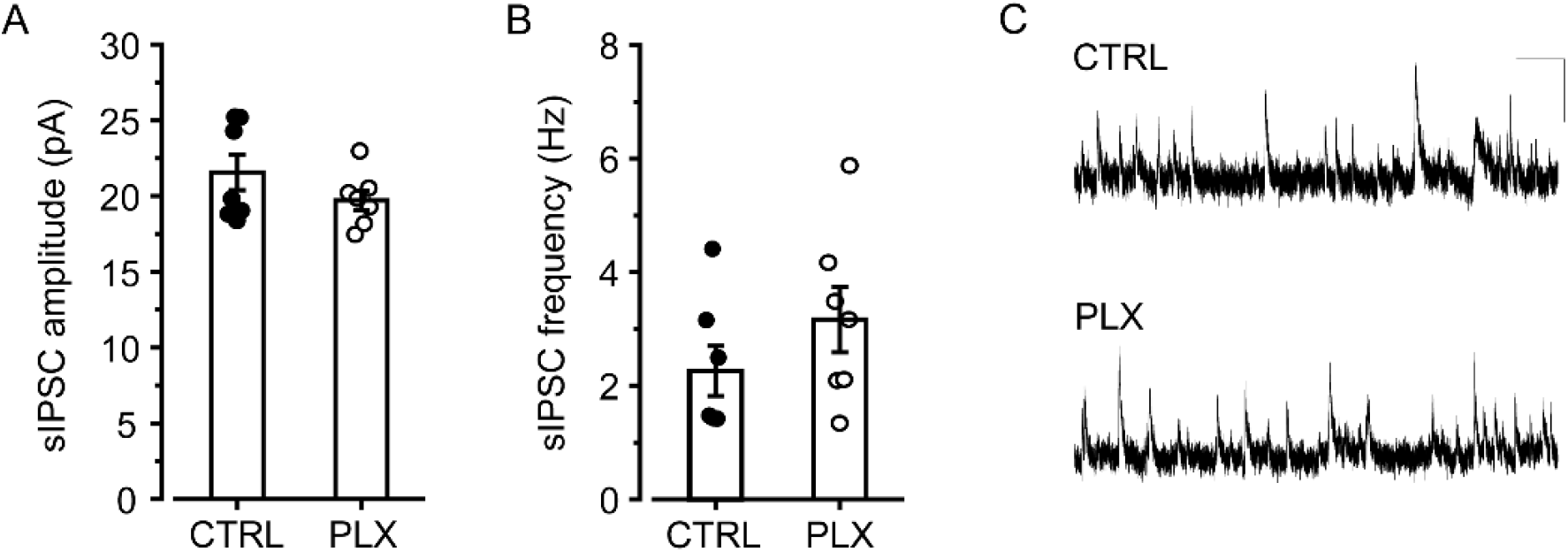
Hippocampal inhibitory transmission is unaltered in PLX-treated mice. A-B) Scatter plots showing no differences in sIPSC mean amplitude and frequency in control (n=7 cells/2 mice) and PLX neurons (n=7 cells/2 mice). In details, for peak current: CTRL 22 ± 1 pA and PLX 19.7 ± 0.7 pA (Mann Whitney test, *p* = 0.4458); for average frequency: CTRL 2.3 ± 0.4 Hz and PLX 3.2 ± 0.6 Hz (Mann Whitney test, *p* = 0.3345). C) Representative sIPSC traces recorded at +10 mV from CA1 pyramidal neurons. Scale bars: 20 pA/1 s. Data shown as Mean ± SEM.

**Appendix Figure S2.**
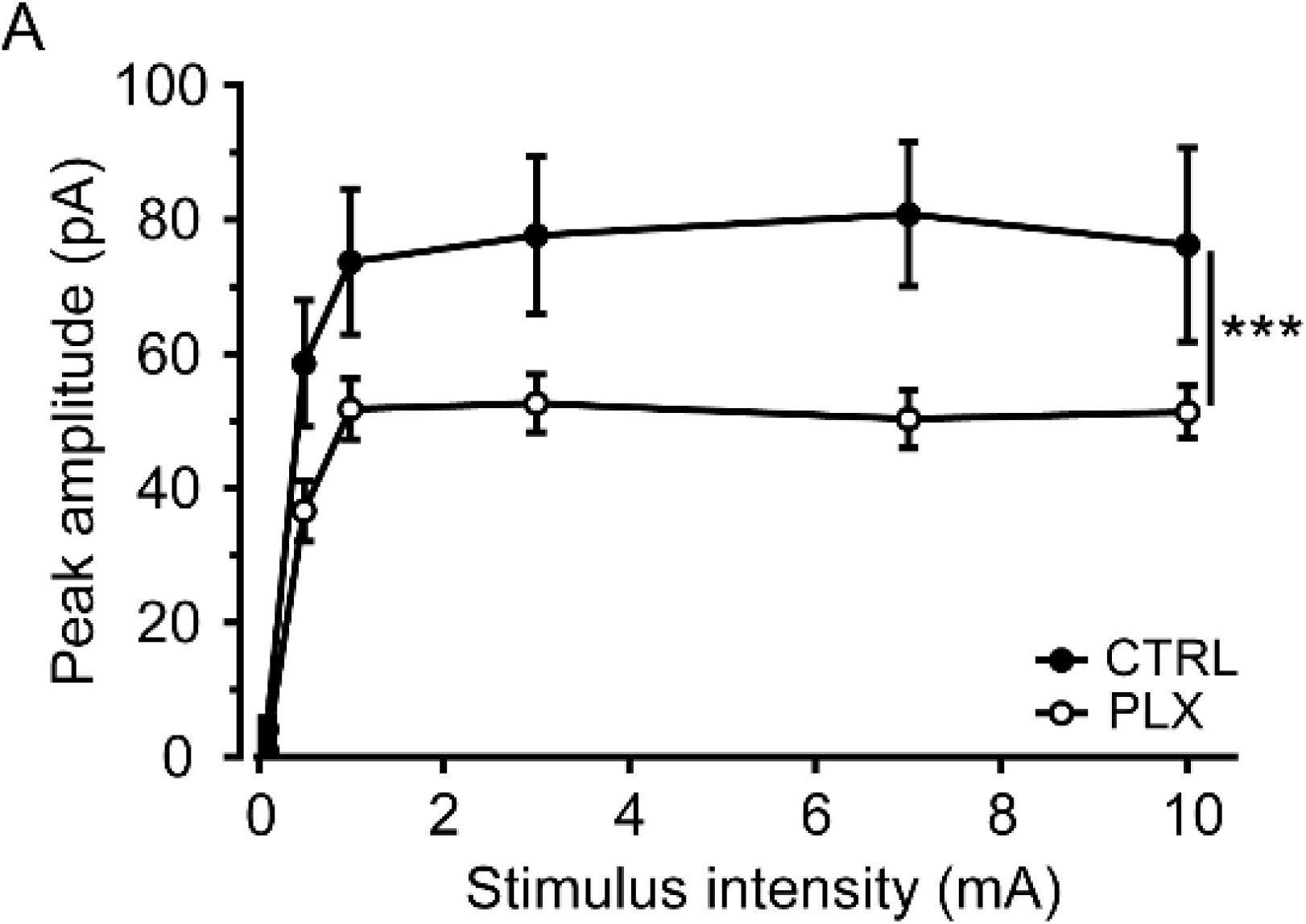
Microglial ablation affects NMDAR-mediated transmission. A) Input-output curve of NMDA component of evoked EPSC. Peak amplitudes recorded at +40 mV in presence of NBQX 10 µM in CA1 neurons from control (n=7 cells/2 mice) and PLX-treated mice (n=9 cells/2 mice). In slices from PLX-treated mice, neurons show significantly lower peak amplitudes compared to control. Two-way ANOVA: treatment F(1,90)=27.53 ***p<.001; stimulation F=(5,90)=26.68 p<.001; interaction F(5,90)=0.95 p>.05. Data shown as Mean ± SEM.

**Appendix Figure S3.**
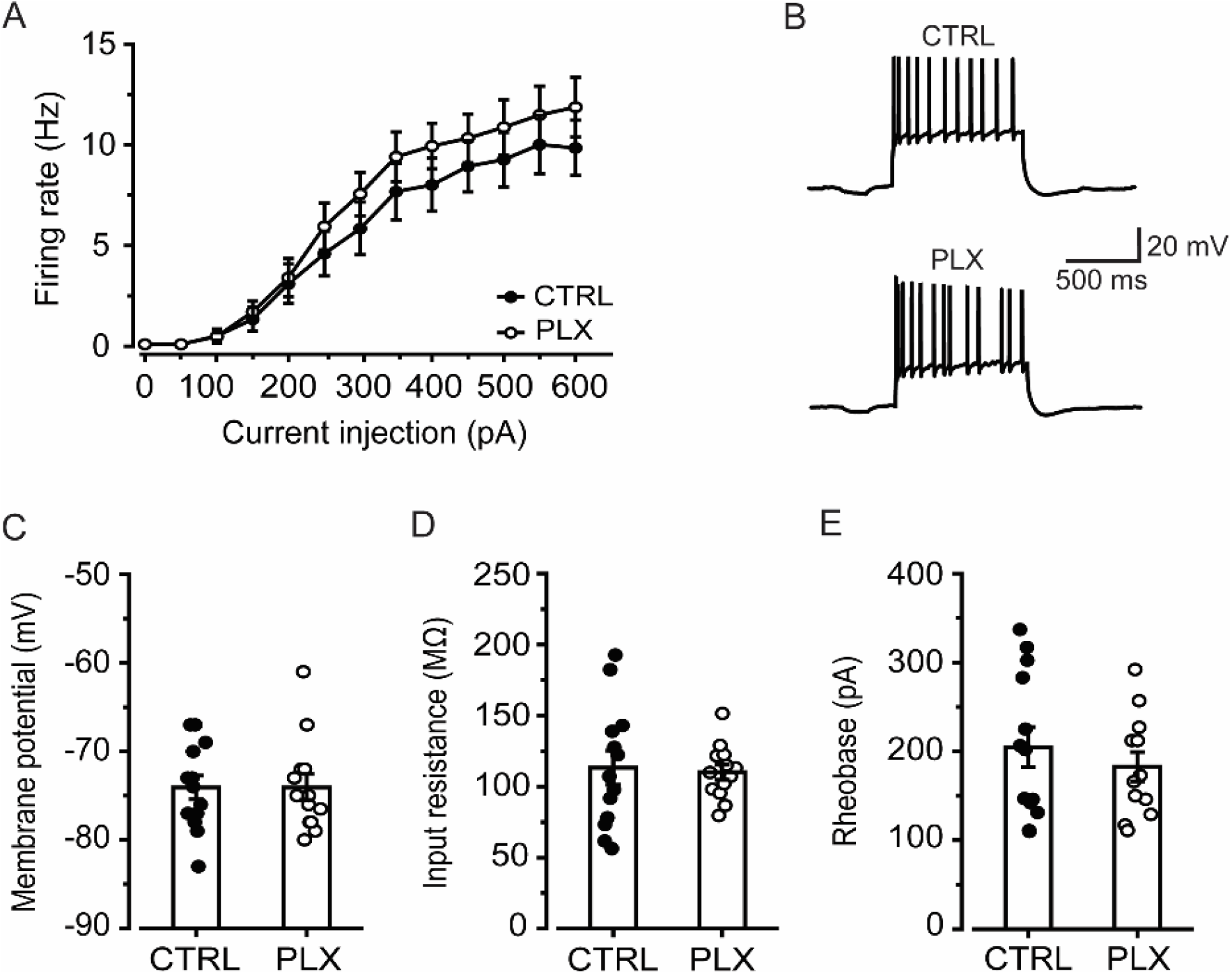
Intrinsic properties and excitability of CA3 pyramidal neurons are unaltered in PLX-treated mice. A) Relationship between injected current (I_inj_) and frequency of evoked action potentials (“*f* – I relationship”) for control (n=12 cells/5 mice) *vs*. PLX-treated animals (n=13 cells/4 mice) depicting lack of difference in evoked excitability. Two-way ANOVA with repeated measures for the I_inj_ range 100 – 600 pA: treatment F(1, 23)=1.017 *p*=0.32; interaction treatment x I_inj_ F(10, 230)=0.4 *p* = 0.94. B) Typical firing evoked in CA3 pyramidal neurons of CTRL or PLX-treated mice in response to steps of injected current (+ 600 pA). C-E) Scatter plots representing lack of alteration of CA3 pyramidal neurons intrinsic properties. In details, membrane potential: CTRL – 74 ± 1 mV and PLX – 74 ± 1 mV (*p* = 0.98; both 13 neurons); membrane input resistance: CTRL 113 ± 12 MOhm and PLX 110 ± 5 MOhm (*p* = 0.8; both 13 neurons); rheobase: CTRL 205 ± 23 pA and PLX 183 ± 17 pA (*p* = 0.44; 13 and 12 neurons, respectively). Data shown as Mean ± SEM.

**Appendix Figure S4.**
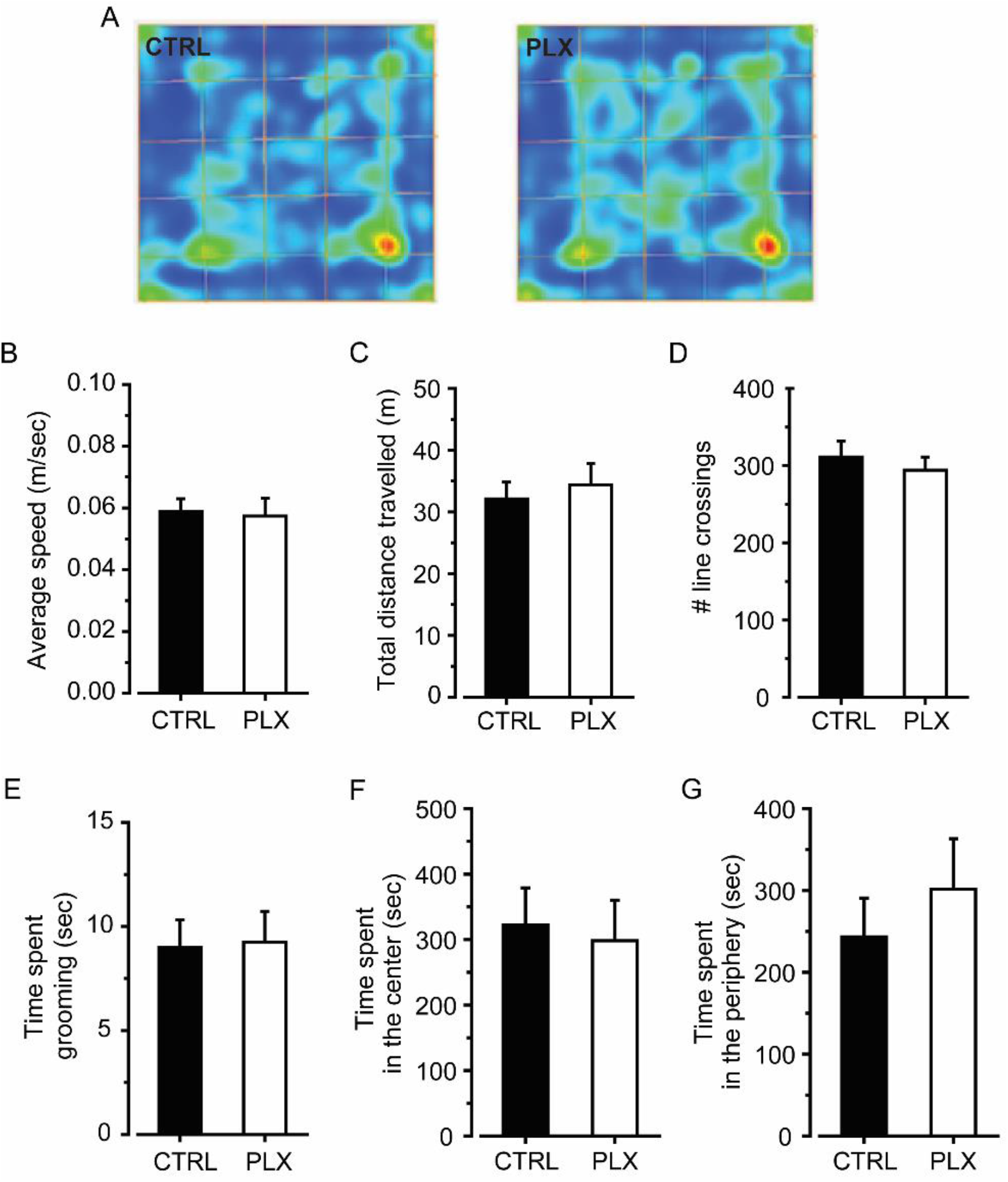
Microglial depletion does not affect locomotor activity and anxiety-like behavior in the Open Field. A) Representative heat maps of locomotor trajectories in the open field arena. B-D) General locomotor activity is not affected by PLX treatment. In particular, no differences were observed in velocity (B), distance travelled (C) and number of crossings (D) in control (n=11) and PLX-treated mice (n=8). t-test t(17)=0.22, p=0.82 (B); t(17)=-0.52, p=0.605 (C); t(17)=0.603, p=0.55 (D). E-G) No differences are observed in grooming behavior (E), time spent in the center (F) and periphery (G) in control (n=11) and PLX-treated mice (n=8). t-test t(17)=-0.12, p=0.901 (E); t(17)=-0.762, p=0.457 (F); t(17)=0.284, p=0.780 (G). Data presented as Mean ± SEM.

